# Arabidopsis Leucine Rich Repeat-Malectin Receptor Kinases in immunity triggered by cellulose and mixed-linked glucan oligosaccharides

**DOI:** 10.1101/2022.08.09.503277

**Authors:** Marina Martín-Dacal, Patricia Fernández-Calvo, Pedro Jiménez-Sandoval, Gemma López, María Garrido-Arandía, Diego Rebaque, Irene del Hierro, Miguel Ángel Torres, Varun Kumar, Diego José Berlanga, Hugo Mélida, Luis F. Pacios, Julia Santiago, Antonio Molina

**Affiliations:** Centro de Biotecnología y Genómica de Plantas, Universidad Politécnica de Madrid (UPM) - Instituto Nacional de Investigación y Tecnología Agraria y Alimentaria (INIA/CSIC), Campus de Montegancedo UPM, 28223-Pozuelo de Alarcón (Madrid), Spain; Departamento de Biotecnología-Biología Vegetal, Escuela Técnica Superior de Ingeniería Agronómica, Alimentaría y de Biosistemas, UPM, 28040-Madrid, Spain; University of Lausanne (UNIL), Biophore Building, Départament de Biologie Moléculaire Végétale (DBMV), UNIL Sorge, CH-1015, Lausanne, Switzerland

**Keywords:** *Arabidopsis thaliana*, cellulose, Mixed-Linked Glucans (MLGs), immunity, oligosaccharides, Pattern Recognition Receptors (PRRs)

## Abstract

Plant immune system perceives through the extracellular ectodomains (ECDs) of Pattern Recognition Receptors (PRRs) a diversity of carbohydrate ligands from plant and microbial cell walls, which activate Pattern-Triggered Immunity (PTI). Among these ligands are oligosaccharides derived from mixed-linked β-1,3/β-1,4-glucans (MLGs, e.g., β-1,4-D-(Glc)_2_-β-1,3-D-Glc, MLG43) and cellulose (e.g., β-1,4-D-(Glc)_3_, CEL3). The mechanisms of perception of carbohydrates by plants are poorly characterized, with the exception of that determining recognition of fungal chitin oligosaccharides (e.g., β-1,4-D(GlcNAc)_6_, CHI6) that involves several PRRs with LysM-ECDs that function as receptor or co-receptors. Here, we describe the isolation and characterization of *Arabidopsis thaliana* mutants *i*mpaired in *<u>g</u>*lycan *<u>p</u>*erception (*igp*), which are defective in PTI activation mediated by MLG43 and CEL3, but not CHI6. *igp1-igp4* are altered in receptor-like kinases [RLKs: AT1G56145 (IGP1), AT1G56130 (IGP2/3), and AT1G56140 (IGP4)] with Leucine-Rich-Repeat (LRR) and Malectin (MAL) domains in their ECDs. *igp4* is a T-DNA insertional, loss of function mutant whereas *igp1* and the allelic *igp2/igp3* harbour point mutations (E906K and G773E, respectively) in their kinase domains, which impact their structure and surface electrostatic potential as revealed by *in silico* structural analyses. Notably, Isothermal Titration Calorimetry assays with purified ECD-RLKs showed that AT1G56145 binds with high affinity CEL3 (Kd = 1.19 ± 0.03 μM) and cellopentaose (Kd = 1.40 ± 0.01 μM), but not MLG43, supporting AT1G56145 function as a plant PRR for cellulose oligosaccharides. Our data suggest that these LRR-MAL RLKs are receptor/co-receptors of a novel mechanism of perception of cellulose and MLG-derived oligosaccharides and PTI activation in *Arabidopsis thaliana*.

**Significance Statement:** New oligosaccharides that trigger plant immunity have been described recently, but the mechanisms of perception of these glycans are unknown. We describe here three *Arabidopsis thaliana* receptor kinases (AT1G56130, AT1G56140, and AT1G56145) with Leucine Rich Repeat (LRR) and Malectin (MAL) domains in their extracellular ectodomains (ECDs), which function as Pattern Recognition Receptors (PRRs) triggering immune response mediated by oligosaccharides from cellulose (β-1,4-glucan) and mixed-linked β-1,3/1,4-glucans (MLGs) of plant and microbial cell walls. The ECD-AT1G56145 binds cellulose oligosaccharides, but not MLGs, supporting its function as a novel receptor of carbohydrate ligands in plants. Our data indicate that these LRR-MAL-PRRs control a complex mechanism of oligosaccharides perception and immune activation that differs from that of fungal chitin oligosaccharides recognition which involves LysM-PRRs.

---

Plants have evolved a very complex immune system that comprises several defence layers and mechanisms for the recognition of pathogens and pests that cooperatively interact to restrict plant infection. One of these layers of defence is known as Pattern-Triggered Immunity (PTI), which is based on the recognition of Damage- and Microbe-Associated Molecular Patterns (DAMPs and MAMPs) derived from plants or microorganisms, respectively, by plasma membrane-resident Pattern Recognition Receptors (1). Upon DAMP/MAMP recognition by ectodomains (ECDs) of PRRs functioning as receptors, additional PRRs are recruited as co-receptors to form PRRs complexes, which trigger the activation of cytoplasmic protein kinase domains (KD) of PRRs that initiate phosphorylation and signalling cascades (1, 2). Among early PTI responses are the increase of cytoplasmic concentration of Ca^+2^ second messenger, the production of reactive oxygen species (ROS) by NADPH-oxidases (e.g., RBOHD), the phosphorylation of Mitogen-activated Protein Kinases (MPKs) and Ca^2+^-Dependent Protein Kinases (CDPKs) and the transcriptional reprogramming of gene expression, that ultimately result in plant restriction of pathogen/pest colonization (2, 3). PTI relevance in plant disease resistance is well-illustrated by the fact that immune responses and restriction of pathogen colonization are compromised in plants defective in PRRs (receptors or co-receptors) involved in the perception of DAMPs or MAMPs of different biochemical nature, including peptides, carbohydrates (oligosaccharides) or fatty acids (1–3).

Many PRR/peptidic DAMP/MAMP pairs triggering PTI responses have been elucidated, like AtPep1 DAMP and bacterial flg22 MAMP peptides which are directly bound by *Arabidopsis thaliana* PEPR1/2 and FLS2 PRRs, respectively (2–4). PRR co-receptors for FLS2 and PEPR1/2 involve members of the SERK family of PRRs, like BAK1, since *bak1* mutant alleles are also impaired in flg22 and AtPep1 perception (2–4). These PRRs are Receptor-Like Kinases (RLKs) with an ECD harbouring Leucine-Rich Repeats (LRR), a Transmembrane domain (TM) and a cytoplasmic serine/threonine KD. RLKs with LRR-ECDs represent about 50% of putative plants PRRs that also comprise Receptor-Like Proteins (RLPs) with ECD and TM but lacking the KD, and Receptor-Proteins (RPs), consisting of an ECD that can be either attached to the plasma membrane by a Glycosylphosphatidylinositol-anchor or to be an extracellular protein (5, 6). *Arabidopsis thaliana* genome has more than 600 genes encoding RLKs/RLPs/RPs members, that in some cases are clustered in the same plant genome loci illustrating their recent evolutionary divergence (1, 6, 7).

In contrast to the extended knowledge of peptidic DAMP/MAMP perception, our understanding of the mechanisms of activation of plant immunity by carbohydrate-based DAMPs/MAMPs is scarce despite carbohydrates being highly abundant in plant and microbial extracellular layers, like cell walls, and that several oligosaccharides that trigger PTI in plants have been identified (8). Moreover, about 50% of RLK/RLP/RP from plant genomes have ECDs that are predicted to bind carbohydrate-based ligands and are grouped in different families: Lysin Motif (LysM), Lectins (G, L and C-lectins), CRinkly-Like (CR4L), Wall-Associated Kinases (WAK), LRR-Malectins, Malectin-LRRs, *Catharanthus roseus* Receptor-Like Kinases 1-like (CrRLK1Ls), and Cysteine-Rich Kinases (CRK/DUF26) (5, 6, 8). Several carbohydrate-based ligands are perceived by the plant immune system such as oligosaccharides from chitin (e.g., chitohexaose (β-1,4-D-N-acetylglucosamine)_6_, CHI6) and β-1,3-glucan of fungal/oomycete cell walls, peptidoglycan from bacterial walls, and oligosaccharides derived from plant cell wall polymers like cellulose (β-1,4-glucan), mix-linked glucans (MLGs; β-1,4/β-1,3-glucans), xyloglucan, mannan, xylan, and homogalacturonan/pectins (oligogalacturonides or OGs) and from other plant glycans, like fructans (9–20).

Cellulose, a linear polymer of β-1,4-glucosyl residues, is present in all plants, most algae, some protists and microbial (bacteria, oomycetes) extracellular matrixes or walls, being the most abundant biomolecule on the earth (21–23). MLGs, consisting of unbranched and unsubstituted chains of β-1,4-glucosyl residues interspersed by β-1,3-linkages, are widely distributed as matrix polysaccharides in cell walls of plants from the Poaceae group (cereals) but have also been reported in *Equisetum* spp. and other vascular plants (24, 25), bryophytes and algae (26, 27), bacteria (28), and fungi and oomycetes (29–31). MLGs derived oligosaccharides (e.g. MLG43 (β-1,4-D-(Glc)_2_-β-1,3-D-Glc), MLG443 (β-1,4-D-(Glc)_3_-β-1,3-D-Glc) and MLG34 (β-1,3-D-Glc-β-1,4-D-(Glc)_2_) are perceived with different degree of specificity by the immune system of several plant species (e.g. *Arabidopsis thaliana*, rice, barley, tomato and pepper) (31–34). Similarly, cellulose-derived oligosaccharides (e.g., β-1,4-D-(Glc)_2_ to β-1,4-D-(Glc)_6,_ cellobiose to cellohexaose, CEL2-CEL6) trigger PTI response in *Arabidopsis thaliana*, rice and other plant species (9, 31, 33–36). MLGs (e.g., MLG43, MLG443) and cellulose-derived oligosaccharides (e.g., CEL2-CEL3) play a critical role as self-alert signals (DAMPs) of plant colonization by pathogens, since they can be released from plant cell walls by the activity of microbial endoglucanases, like cellulases, secreted during plant colonization (33, 37). MLGs can be also perceived as MAMPs by those plant species that do not contain MLGs in their cell walls, such as *Arabidopsis thaliana* and Solanaceae species (31, 32).

Our knowledge of the mechanisms of oligosaccharides perception by the mammal immune system through carbohydrate-recognition domains (CRDs) is quite large (8, 38, 39). In contrast, our understanding of the structural basis of perception of oligosaccharides by plants PRR is limited and mainly restricted to PRRs of the LysM family (e.g., CERK1 (Chitin Elicitor Receptor Kinase 1), LYK4 and LYK5), which harbour promiscuous ECDs involved in the activation of PTI mediated by structural diverse ligands like chitin derived oligosaccharides (e.g., CHI6-CHI8), β-1,3-glucan (laminarihexaose, LAM6) and MLGs (MLG43/MLG34/MLG443) (15, 19, 31, 33, 40–44). Immune responses triggered by CHI6, LAM6 and MLG43 are partially impaired in *cerk1* single and *lyk4 lyk5* double mutants (15, 31, 43-46). However, a direct binding of LAM6 and MLG43 to *Arabidopsis thaliana* CERK1 ECD was discarded based on *in silico* structural molecular dynamics simulations and Isothermal Titration Calorimetry (ITC) binding assays performed with purified ECD-CERK1 and LAM6 (6). These data suggest that CERK1/LYK4/LYK5 might function as a co-receptor or be part of LAM6 perception system. Notably, rice OsCERK1 seems to bind MLG43 and MLG443, whereas OsCeBIP, the rice chitin receptor, is required for these ligands’ perception, but not for their direct binding (33). In addition to LysM-PRRs, Arabidopsis WAK receptors (e.g., WAK1 and WAK2) have been described as PRRs required for the perception of OGs, and the CrRLK1L member FERONIA (FER), which has two MAL domains in its ECD, has been described to bind homogalacturonans, but the structural bases of these bindings have not been elucidated yet (47, 48).

To identify novel PRRs involved in the recognition of glycans DAMPs/MAMPs and to characterize novel mechanisms of PTI activation by oligosaccharides in plants, we designed genetic screenings in *Arabidopsis thaliana* to identify mutants *i*mpaired in *<u>g</u>*lycan *<u>p</u>*erception (*igp*). We selected several mutants (*igp1-igp9*) impaired in the perception of MLG43 and CEL3, but not CHI6, since cytoplasmic Ca^2+^ elevations and additional PTI responses were partially or fully impaired in *igp1-igp9* upon treatment with either MLG43 or CEL3. We show here that *IGP1, IGP2/IGP3,* and *IGP4* encode three previously uncharacterized RLK members (AT1G56145, AT1G56140, and AT1G56130, respectively) with LRR-MAL in their ECDs, which are required for perception of MLG43 and CEL3 and PTI activation. We also demonstrate that the ECD of AT1G56145 binds directly to CEL3 and CEL5 with high affinity, further supporting its function as a plant PRR receptor of cellulose-derived oligosaccharides. These results expand our knowledge of oligosaccharide perception and immune activation in plants.

## Results

### Identification and isolation of Arabidopsis *igp1-igp9* mutants

We generated an ethyl methanesulphonate (EMS) mutagenized population of the *Arabidopsis thaliana* Aequorin-based Ca^2+^sensor line (Col-0^AEQ^: (15, 49, 50) to perform a genetic screening aiming to isolate mutants *i*mpaired in *g*lycans *p*erception (*igp*). This screening was based on the identification of mutant lines (*igp)* with defective activation of cytoplasmic Ca^2+^ burst upon application of oligosaccharides in comparison to Col-0^AEQ^ (*SI Appendix*, Fig. S1*A*). Eight-day-old seedlings (about 6,400 individuals) of 13 M2 EMS-mutagenized Col-0^AEQ^ families and of Col-0^AEQ^ control line were grown in microtiter plates, treated with 100 μM of MLG43 and triggered Ca^2+^ burst determined in a luminometer (*SI Appendix*, Fig. S1*A*). Several mutants (*igp1* ^AEQ^*-igp3*^AEQ^ and *igp5*^AEQ^*-igp9*^AEQ^) that showed weaker Ca^2+^ burst than that of Col-0^AEQ^ plants were selected for further genetic characterization (backcrosses and allelism tests: Fig. 1*A* and *SI Appendix*, Fig. S2). The specificity of MLG43 perception impairment in *igp* was assessed by determining Ca^2+^ burst upon their treatment with fungal MAMP CHI6 (50 μM). This analysis revealed that Ca^2+^ bursts were similar in *igp* mutants and Col-0^AEQ^ plants upon CHI6 treatment, in contrast to the impaired response of *cerk1*^AEQ^ line, as reported (Figure 1*B* and *SI Appendix*, Fig. S2;15), indicating that the mechanism of perception of CHI6 and MLG43 in *Arabidopsis thaliana* might not be identical, as suggested previously (31). To further discard that Ca^2+^ burst reduction observed in MLG43-treated *igp* mutants might be due to mutations in the *Aequorin* transgene, we determined the sequence of *Aequorin* gene in *igp1*^AEQ^*-igp3*^AEQ^ lines, and mutations were not found (*SI Appendix* Dataset 1). Moreover, we also performed Ca^2+^ discharges in *igp1*^AEQ^*-igp3*^AEQ^ and Col-0^AEQ^ plants to determine total Ca^2+^ in seedling cells and no differences were observed in endogenous Ca^2+^ levels between the mutants and Col-0^AEQ^ seedlings (*SI Appendix*, Fig. S1*B*), further indicating that the observed lower Ca^2+^ bursts in MLG43 treated *igp1*^AEQ^*-igp3*^AEQ^ seedlings were due to a defective perception of MLG43.

**Figure 1.**
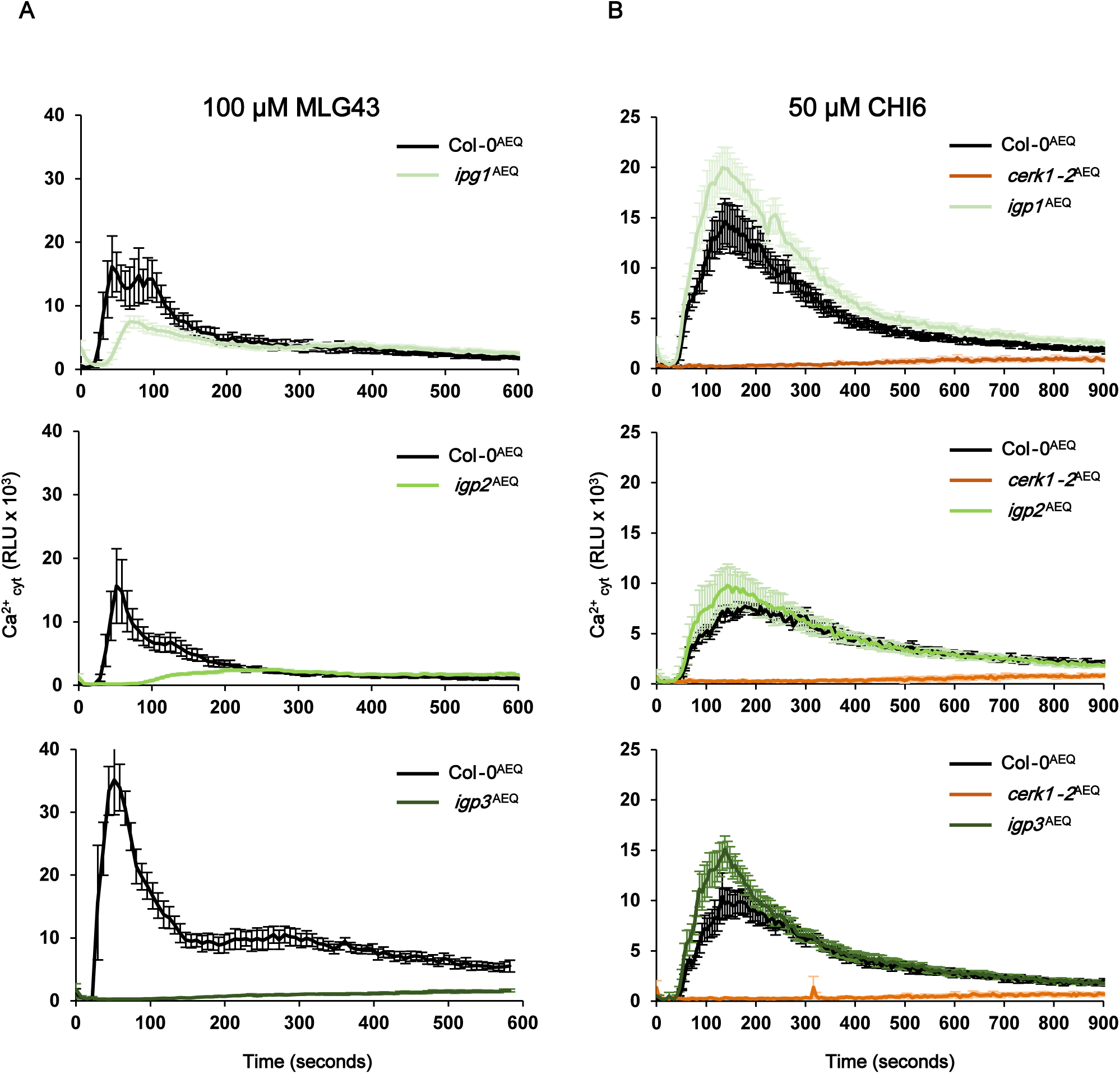
Identification of *Arabidopsis thaliana* mutants *i*mpaired in *g*lycans *p*erception (*igp*). Cytoplasmic calcium burst upon application of (*A*) 100 µM of MLG43 and (*B*) 50 µM CHI6) was measured as relative luminescence units (RLU) over the time in 8-day-old seedlings of Col-0^AEQ^ and *igp*^AEQ^ mutants. *cerk1-2*^AEQ^ line, impaired in CHI6 perception, was included for comparison. Data represent the mean ± standard error (n = 4 in the case of Col-0^AEQ^ and *cerk1-2*^AEQ^; and n = 8 in the case of *igp*^AEQ^ mutants). Data are from one of the four experiments performed that gave similar results.

*igp1*^AEQ^, *igp2*^AEQ^ and *igp3*^AEQ^ mutants harbour recessive mutations (Chi-test: 0,7 > *P* > 0,5; 0,5 > *P* > 0,3 and 0,8 > *P* > 0,7, respectively; *SI Appendix,* Dataset 2) and *igp2*^AEQ^ and *igp3*^AEQ^ were allelic (Chi-test: *P* > 0,95; *SI Appendix,* Dataset 2, Fig. S1*C*). In the progeny of F2 backcrosses of *igp1*^AEQ^*, igp2*^AEQ^, and *igp3*^AEQ^ with Col-0^AEQ^ we selected 50 plants impaired in MLG43 perception, bulk genomic DNA was extracted from these plants, and Col-0^AEQ^ parental lines and DNA was sequenced and assembled to identify putative mutations (frequency higher than 0.99 in alignments: *SI Appendix* Dataset 3). We found that *igp1*^AEQ^ has a point mutation in the *AT1G56145* gene, encoding an RLK with an LRR-MAL ECD, that resulted in E906K amino acid change in its KD (Fig. 2*A*). On the other hand, *igp2*^AEQ^ *and igp3*^AEQ^ share the same point mutation in the *AT1G56130* gene, encoding an additional LRR-MAL RLK, that resulted in G773E amino acid change in its KD (Fig. 2*A*). Notably, these two LRR-MAL RLKs are in a genomic cluster with two additional LRR-MAL RLK encoding genes, *AT1G56120* and *AT1G56140*, which are also members of the LRR-MAL RLK family comprising at least 12 genes in the *Arabidopsis thaliana* genome (6; *SI Appendix*, Fig. S3*A*), with a few characterized members, like *RFK1* (*AT1G29720*) that promotes compatible pollen grain hydration and pollen tube growth (51). Phylogenetic analyses indicated that these four genes (*AT1G56120/AT1G56130/AT1G56140/AT1G56145)* form part of a specific clade of the LRR-MAL RLK family, that differs from RLKs with MAL-LRR ECDs (*SI Appendix*, Fig. S3*A*).

**Figure 2.**
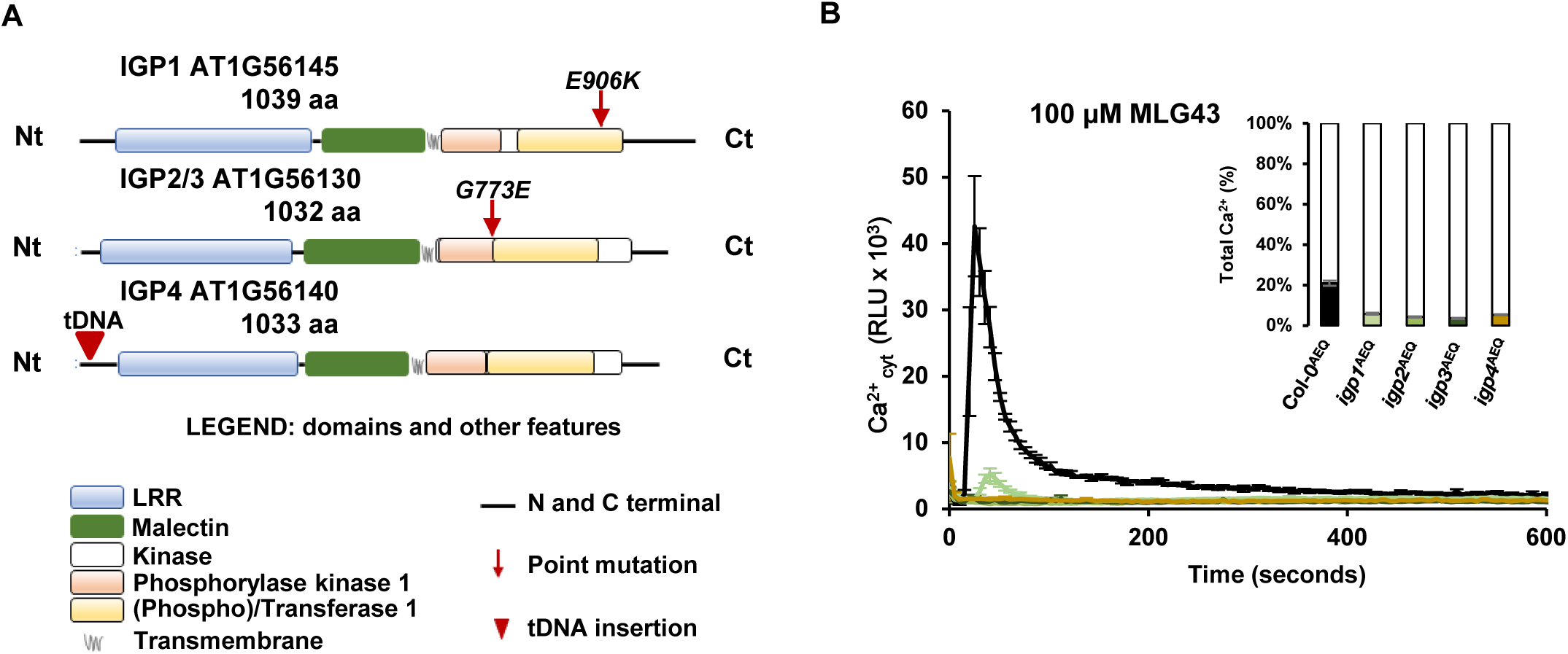
Identification of casual mutations in *igp1-igp4* mutants. (*A*) Representation of protein domains of AT1G56130, AT1G56140 and AT1G56145 LRR-MAL RLKs: LRR (blue), MAL (green), phosphorylase kinase (orange) and phophotransferase (yellow) of KD, regions connecting such domains and N- and C-terminal domains (black lines), and TM (grey). Red arrows indicate the position of the mutations in the coding regions of *igp1*^AEQ^, *igp2*^AEQ^ and *igp3*^AEQ^, and the red triangle the insertion of the T-DNA sequence in *igp4*. Whole genome sequencing and single polymorphism analysis led to the identification of a C to T changes in DNA sequences of *AT1G56130* and *AT1G56145* genes (*SI Appendix Data Set 1*), that led to G to E, and E to K amino acid changes in the indicated amino acids of RLK KDs. (*B*) Calcium burst measured as relative luminescence units (RLU) over the time in 8-day-old Col-0^AEQ^, *igp1*^AEQ^, *igp2*^AEQ^, *igp3*^AEQ^, and *igp4*/*at1g56140*^AEQ^ seedlings upon treatment with 100 µM MLG43. Data represent the mean ± standard error (n=4 in the case of Col-0^AEQ^ and n=8 in the case of *igp*^AEQ^ mutants). After recording the elicitor induced Ca^2+^ kinetics, the total Ca^2+^ was discharged by the addition of 1mM CaCl_2_ to the wells. The kinetic areas after both treatments were integrated and their values used for the calculation of the total Ca^2+^ % induced by the treatment compared with the total calcium discharge (graph at the top right). This is one of the three experiments performed that gave similar results.

We tested by qRT-PCR the expression of these 4 LRR-MAL RLK encoding genes in seedlings treated with MLG43, CHI6 and other glucans (e.g., CEL3) and we found that the expression of *AT1G56140,* but not that of *AT1G56120*, *AT1G56130 and AT1G56145* was up-regulated upon treatment with these oligosaccharides (*SI Appendix*, Fig. S3*B*). We then selected T-DNA insertional mutants of *AT1G56120* and *AT1G56140* genes and tested their perception of MLG43 and CHI6 by determining by qRT-PCR the upregulation of PTI marker genes *WRK53* and *CYP81F2* (15, 31) in treated seedlings of these mutants and Col-0 wild-type plants (*SI Appendix*, Fig. S3*C*). In the *at1g56140* knock-out mutant a reduced upregulation of *WRK53* and *CYP81F2* expression upon MLG43 treatment was clearly observed in comparison to Col-0 lines, whereas in *at1g56120* mutant the upregulation of PTI marker genes upon MLG43 or CHI6 treatments was very similar to that of Col-0 plants (*SI Appendix*, Fig. S3*C*). In both mutants, up-regulation of *WRK53* and *CYP81F2* expression upon CHI6 treatment was similar to that of wild-type plants (*SI Appendix*, Fig. S3*C*). To further validate the phenotype of the *at1g56140* mutant, it was crossed with Col-0^AEQ^ and the homozygous *at1g56140*^AEQ^ line generated was tested for Ca^2+^ burst upon MLG43 and CHI6 treatment. We found that *at1g56140*^AEQ^ was impaired in MLG43, but not CHI6, perception, similarly to *igp1*^AEQ^*-igp3*^AEQ^, and that this defective response was not due to alteration in Ca^2+^ endogenous levels, as revealed by Ca^2+^ discharge experiments, and accordingly the mutant was named *igp4^AEQ^* (Fig. 2*B*). These results indicate that *AT1G56145* (*IGP1*), *AT1G56130* (*IGP2/IGP3*) and *AT1G56140* (*IGP4*) LRR-MAL RLKs are required for MLG43, but not CHI6, perception, and PTI activation. We assessed the impact of the *igp1-igp4* mutations on plant developmental phenotypes and no significant differences were observed in the rosette and silique size and morphology of *igp1*^AEQ^ and *igp4*^AEQ^ plants in comparison to Col-0^AEQ^ or *cerk1-2*^AEQ^, whereas *igp2*^AEQ^ and *igp3*^AEQ^ plants have rosettes and siliques slightly smaller than those of Col-0 plants or *cerk1-2*^AEQ^ (*SI Appendix*, Fig. S4*A-C*).

### LRR-MAL RLKs are also required for the perception of additional cellulose and MLG derived oligosaccharides

To determine the specificity of PTI responses mediated by this group of LRR-MAL RLKs, we treated *igp1^AEQ^*, *igp3^AEQ^*and *igp4^AEQ^* and Col-0^AEQ^ seedlings with different DAMPs (i.e., carbohydrates ligands CEL3 and OGs (DP10-12), and AtPep1) and the MAMP flg22, and Ca^2+^ bursts were measured. Remarkably, the three *igp*^AEQ^ lines were almost fully impaired in CEL3-mediated Ca^+2^ burst activation, suggesting that they are also required for the perception of cellulose-derived oligosaccharides (Fig. 3*A,B*). In contrast, the elevation of cytoplasmatic Ca^2+^ induced upon flg22, OGs and AtPep1 treatments in the mutants were similar to those observed in Col-0^AEQ^ (Fig. 3*C-E*). We also tested the response of *igp5*^AEQ^*-igp9*^AEQ^ mutants to CEL3 and we found that they were also impaired in CEL3 perception (*SI Appendix*, Fig. S5), further supporting that the mechanisms of perception of MLG43 and CEL3 in *Arabidopsis thaliana* share some components, like the three PRRs with LRR-MAL ECDs. Since additional cellulose and MLGs derived oligosaccharides, like cellobiose (CEL2), cellotetraose (CEL4), cellopentaose (CEL5) and MLG34 have been described to trigger PTI in *Arabidopsis thaliana* (31, 35, 36), we also determined Ca^2+^ burst activated by these glycans in *igp1*^AEQ^, *igp3*^AEQ^ and *igp4*^AEQ^ and Col-0^AEQ^. As shown in *SI Appendix*, Fig. S6, *igp1*^AEQ^, *igp3*^AEQ^ and *igp4*^AEQ^ showed reduced Ca^2+^ influxes in comparison to Col-0^AEQ^ upon treatment with CEL4, CEL5 and MLG34, indicating that the three LRR-MAL RLKs are required for perception of these cellulose and MLGs derived oligosaccharides. Ca^2+^ influxes triggered by CEL2 were very low in comparison to the rest of the active ligands tested, suggesting that this disaccharide has low immunogenic activity in *Arabidopsis thaliana* and that CEL3 is the minimal, active cellulose-derived oligosaccharide putatively recognized by these RLKs (*SI Appendix*, Fig. S6).

**Figure 3.**
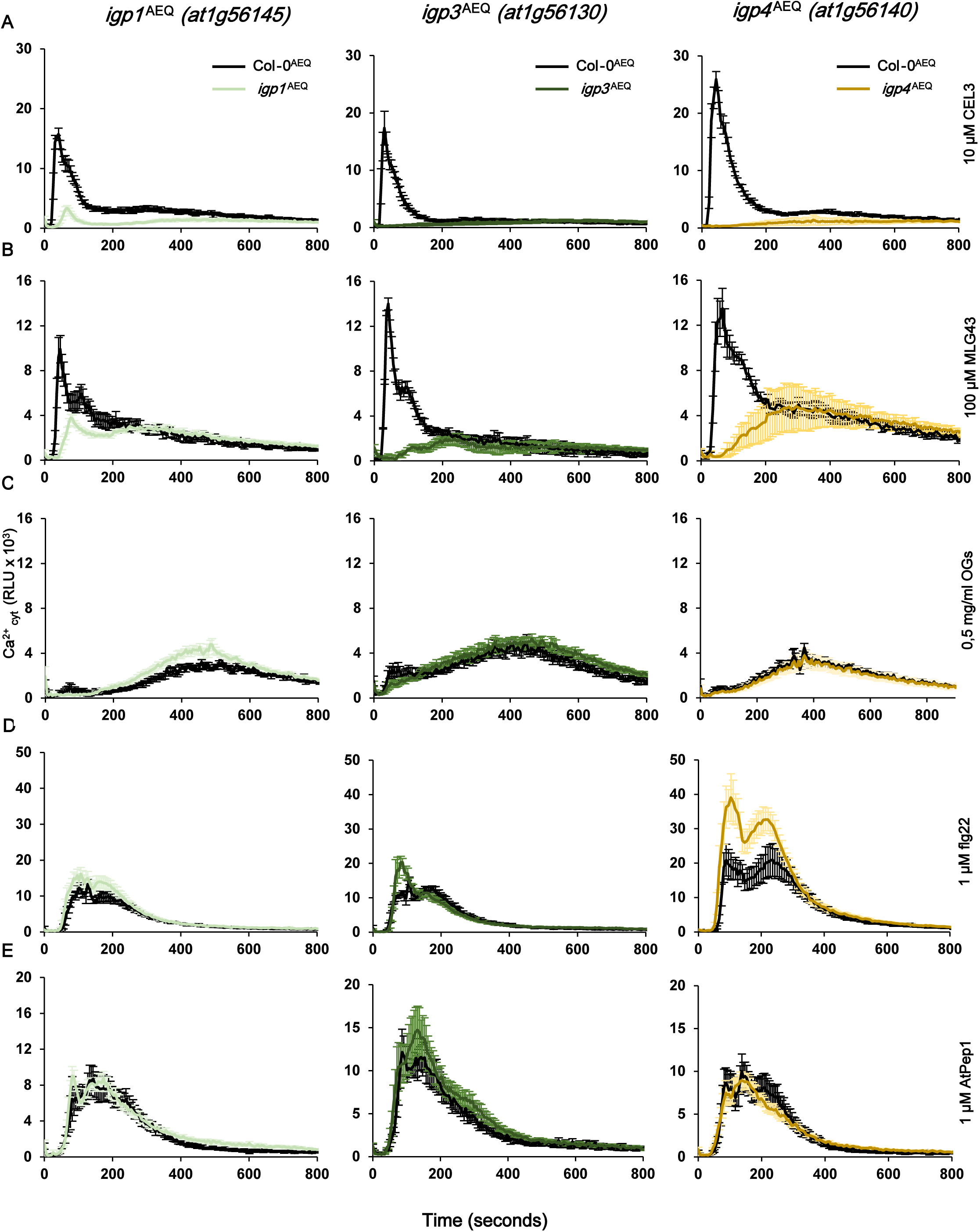
Cytoplasmic calcium burst triggered by CEL3 is impaired in *igp*^AEQ^ mutants. Ca^2+^ burst measured as relative luminescence units (RLU) over the time in 8-day-old Col-0^AEQ^ in *igp1*^AEQ^, *igp3*^AEQ^ and *igp4*^AEQ^ seedlings after treatment with (*A*) 10 µM CEL3, (*B*) 100 µM MLG43, (*C*) 0,5 mg/ml OGs, (*D*) 1 µM flg22, and (*E*) 1 µM AtPep1. Data represent the mean ± standard error (n=4 in the case of Col-0^AEQ^ and n=8 in the case of *igp*^AEQ^ mutants). Data are from one of the three experiments performed that gave similar results.

To further validate whether the mechanism of perception of cellulose and MLGs oligosaccharides in *Arabidopsis thaliana* share some PRRs and signalling components, we performed cross-elicitation experiments by treating 8-day-old Col-0^AEQ^ seedlings first with MLG43 or CEL3, and a few minutes later with either CEL3 or MLG43. As shown in *SI Appendix* Fig. S7, Ca^2+^ influxes were not observed upon treatment of Col-0^AEQ^ seedling with MLG43+CEL3, similarly to what was observed in CEL3+CEL3 and MLG43+MLG43 treatments, indicating that PRRs or components involved in the perception and downstream activation of Ca^2+^ influxes mediated by MLG43 and CEL3 are shared in *Arabidopsis thaliana*.

To further confirm the role of this group of LRR-MAL receptors as potential PRRs for cellulose and MLG oligosaccharides perception in *Arabidopsis thaliana*, we analysed the activation of additional immunity hallmarks in *igp1*^AEQ^*, igp3*^AEQ^ and *igp4*^AEQ^ and Col-0 wild-type plants upon treatment with MLG43, CEL3 and CHI6. We first monitored ROS burst in these genotypes and *rbohD* line, which is impaired in DAMP/MAMP-triggered ROS production (52). Upon treatment with MLG43, ROS burst was partially impaired in *igp* mutants in comparison to Col-0 plants, whereas in CEL3-treated *igp* mutants ROS burst was significantly reduced to a higher extent (Fig. 4*A,B*). In both cases, the reduction of ROS production was not as noticeable as in the case of *rbohD* (Fig. 4*A,B*). Notably, ROS burst in *igp* mutants upon CHI6 and flg22 treatment was not altered in comparison to Col-0 plants (*SI Appendix*, Fig. S8*A,B*), as observed for Ca^+2^ burst (Fig. 3). Next, we tested the phosphorylation of protein kinases (MPK3/MPK6/MPK4/MPK11) by Western-blot and we found that MPK3- and MPK6-phosphorylation triggered by MLG43 and CEL3 was reduced in *igp1*^AEQ^*, igp3*^AEQ^ and *igp4*^AEQ^ in comparison to Col-0 plants, whereas MPK3/MPK6 phosphorylation in CHI6 treated *igp* mutants was similar to that of Col-0 plants (Fig. 4*B*). MPK4/11-phosphorylation was almost not-detectable in either CEL3-treated or MLG43-treated plants (Fig. 4*B*), as reported previously (31). Last, we studied by qRT-PCR the expression of two PTI-marker genes up-regulated by CHI6 and MLG43 (*WRKY53* and *CYP81F2;* (15, 31)) in seedlings 30 minutes after treatment with these oligosaccharides. We found that up-regulation of *WRKY53* and *CYP81F2* in MLG43- and CEL3-treated *igp1*^AEQ^*, igp3*^AEQ^ and *igp4* ^AEQ^ seedlings was weaker than in Col-0 plants (Fig. 4*C*). In contrast, in CHI6-treated *igp* mutants, transcriptional up-regulation of these genes was similar to that of Col-0 plants (Fig. 4*C*).

**Figure 4.**
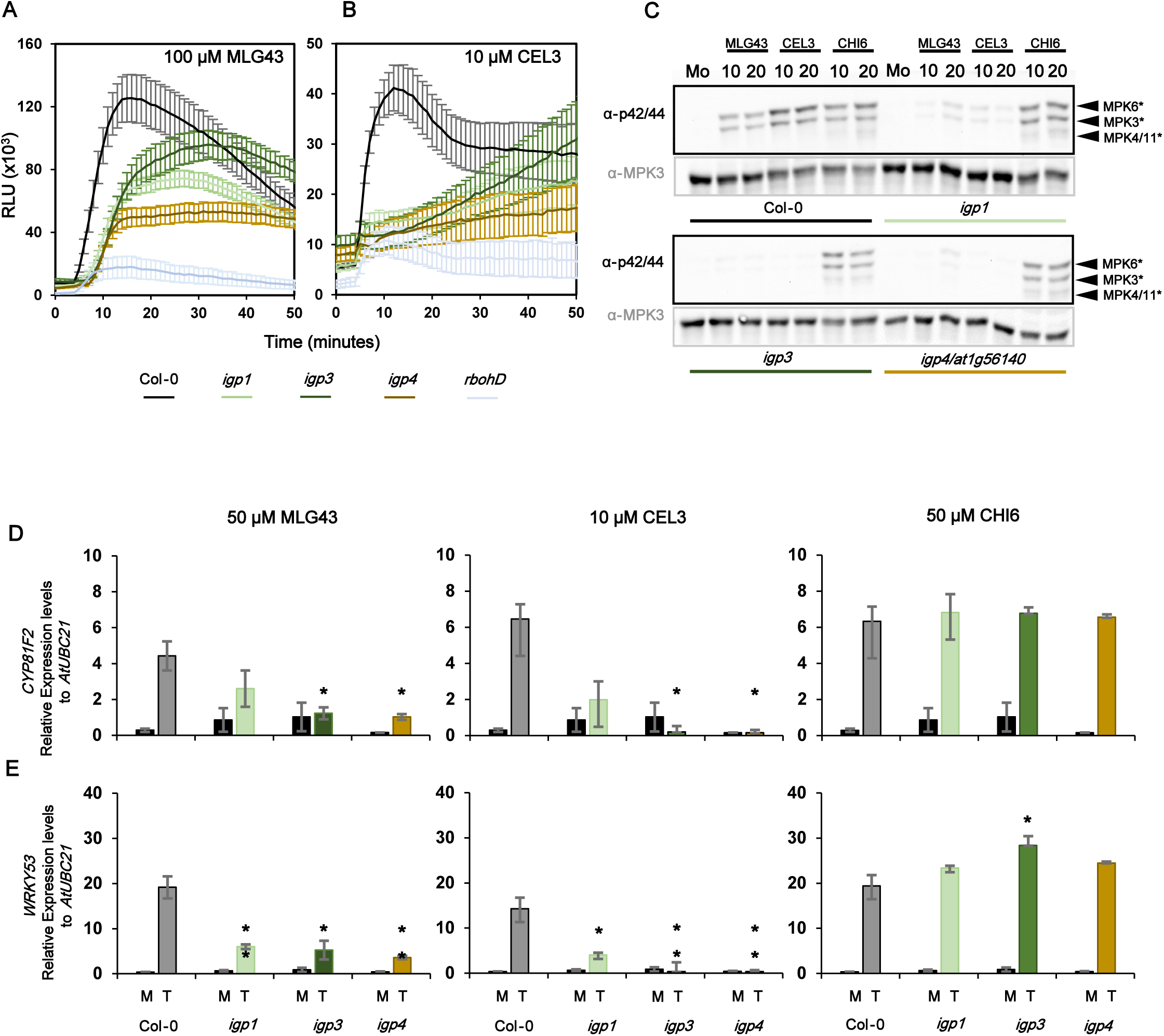
Activation of PTI hallmarks by MLG43, CEL3 and CHI6 in *igp*^AEQ^ mutants. *(A,B)* Reactive Oxygen Species (ROS) production was monitored as H_2_O_2_ production over a period of 50 minutes by luminol-assays and measured as relative luminescence units (RLU) in the indicated genotypes. Oligosaccharides were added 5 minutes after the transfer of the plate with seedling and luminol reagents to the luminometer: (*A*) 100 μM MLG43 and (*B*) 10 μM CEL3. Data represent mean ± standard error (n = 8). Comparison to Col-0 assessed by Student t-test (n = 24) at the time of Col-0 peak shows statistically significant differences for *igp1*^AEQ^ and *igp3*^AEQ^ (*P* < 0.01) and *igp4*^AEQ^ (*P* < 0.001) in response to MLG43. Likewise, all three mutants show statistical differences at *P* < 0.001 compared to Col-0 in response to CEL3. *rbohD* plants impaired in ROS production showed statistically significant differences (*P* < 0.001) with both treatments. These results are from one representative experiment out of the three performed that gave similar results. (*C*) Mitogen-activated protein kinases (MAPK) phosphorylation was analyzed in seedlings of Col-0 and *igp1*^AEQ^, *igp3*^AEQ^ and *igp4*^AEQ^ treated with 100 µM MLG43, 10 µM CEL3, 50 µM CHI6 or water (mock). Western-Blot using anti-pTEpY antibody (anti p42/44) for phosphorylated MAPK moieties was performed with samples harvested at different time points (0, 10 and 20 minutes). Black arrows indicate the position of phosphorylated MPK6 (top), MPK3 (middle) and MPK4/11 (bottom). Anti-MPK3 was used as total protein control to show the loading of each gel. These results are from one representative experiment out of the two performed that gave similar results. (*D)-(E*) Quantitative RT-PCR analysis in 12-day-old seedlings of the indicated genotypes. Expression levels of immune marker genes (*D*) *CYP81F2* and (*E*) *WRKY53*, relative to the *Arabidopsis thaliana* housekeeping gene *UBC21* (*AT5G25769*) at 30 min after mock treatment or application of the oligosaccharides are shown. Data represent mean ± standard error of three technical replicates out of three independent biological replicates (n=3). Statistically significant differences between MLG43, CEL3 or CHI6 treated *igp*^AEQ^ *versus* treated Col-0 according to Student’s t-test (\**P* < 0.05, ** 0.01< *P* > 0.001, *** *P* < 0.001).

Since Arabidopsis and rice perception of MLGs have been described to involve LysM-PRRs (e.g., CERK1, LYK4 and LYK5 in *Arabidopsis thaliana*), we determined MPK3/MPK6 phosphorylation, ROS production and gene expression upon treatment with MLG43 or CEL3 in seedlings of Col-0, *cerk1-2* and *cerk1-2lyk4lyk5* triple mutant, which was generated in this work (*SI Appendix*, Fig. S4*A-C*). ROS kinetics and burst in plants treated with MLG43 were slightly lower in *cerk1-2 and cerk1-2lyk4lyk5* genotypes than in Col-0, whereas it was not impaired in *cerk1-2 and cerk1-2lyk4lyk5* genotypes treated with CEL3 (*SI Appendix*, Fig. S9*A*). Moreover, a slight reduction of the phosphorylation of MPK3/MPK6 in these mutants compared to Col-0 was observed upon MLG43 and CEL3 treatment, but it was weaker than that observed in LysM-PRR mutants treated with CHI6 (*SI Appendix*, Fig. S9*B*). MLG43-mediated up-regulation of *WRKY53* and *CYP81F2* was partially affected in *cerk1-2* and *cerk1-2lyk4lyk5,* whereas it was not altered upon CEL3 treatment (*SI Appendix*, Fig. S9*C*). Together these PTI hallmark analyses further support the function of the three LRR-MAL RLKs identified as PRRs involved in the perception of cellulose (either as receptors or co-receptors), and in lower extent, of MLGs derived oligosaccharides. These data also indicate that MLG43 perception involves CERK1/LYK4/LYK5 LysM-PRRs, as described previously for MLGs in *Arabidopsis thaliana* and rice (31, 33), whereas these LysM-PRRs have almost no contribution in CEL3 perception.

### Model structures of AT1G56130, AT1G56140 and AT1G56145 RLKs point to their function as PRRs

Malectin domains (MAL) like those present in ECDs of the LRR-MAL RLK family have been previously described in animals to bind short glycans based on the NMR structure of malectin from *Xenopus laevis* in complexes with maltose and nigerose (53). MAL domain is present in at least 2 additional families of plant RLKs (MAL-LRRs, and CrRLK1Ls), and the ECDs of two CrRLK1Ls members (ANXUR1 (ANX1) and ANXUR2 (ANX2)) which have been crystallized, but no oligosaccharide ligands were identified in ITC binding experiments (54). We performed phylogenetic analyses of MAL domains of *Arabidopsis thaliana* LRR-MAL RLK members (represented by AT1G56145) and determined the conservation of MAL domain in plants, finding that it is highly conserved in cruciferous and other dicots species, but orthologs seems to be absent in grasses (*SI Appendix*, Fig. S10). On the other hand, the MAL domain of AT1G56130/AT1G56140/AT1G56145 were found to be evolutionary divergent from ANX1 and ANX2 and *Xenopus* sp. MAL domains (*SI Appendix*, Fig. S10).

We then performed *in silico* analyses on the predicted AlphaFold LRR-MAL RLK model structures to understand the impact of *igp1* and *igp2/igp3* point mutations in the KD of AT1G56130 and AT1G56145. AlphaFold predicts LRR, MAL and KD domains of the three RLKs with confidence *very high* (> 90%) for LRR and KD, *high* for MAL (70–90) and *low* (50-70%) or *very low* (< 50%) for transmembrane (TM) and both N- and C-terminal regions, respectively (*SI Appendix*, Fig. S11A-C). In AlphaFold the spatial arrangement of all these domains does not properly describe the expected ECD (LRR+MAL)-TM-intracellular KD organization (*SI Appendix*, Fig. S11*D)*. Therefore, we modified AlphaFold structures to obtain a proper domains organization (see Material and Methods). The new models (Fig. 5*A*) provide a complete picture of the proteins to be used as initial geometries for further computational studies (e.g., docking, and molecular dynamics simulations). To assess the models of MAL domains (which have lower confidence than LRR and KD domains), we selected for MAL domain structural modeling some of the few entries in PDB (5 plant RLKs from CrRLK1Ls involved in peptide perception and 9 human and bacteria proteins), including the NMR structures of malectin from *X. laevis* in complex with maltose (PDB code 2KR2; (53)), *X. laevis* malectin (an apo-form and a complex with nigerose: (55)), and the crystal structure of tandem malectin-like domains in the ANX1/ANX2 ECDs from *Arabidopsis thaliana* (PDB code 6FIG; (54)). MAL domains in AlphaFold models of AT1G56130 (202 residues), AT1G56140 (198 residues), and AT1G56145 (196 residues) are extremely similar: the three pair Combinatorial Extension (CE) structural superpositions included 192 residues with Root-Mean Square Deviation (RMSD) values 0.730-0.790 Å (Fig. 5*B*). As for MAL experimental structures and taking AT1G56140 as a reference, the CE superposition with 2KR2 (179 residues) and 6FIG (224 residues) included 146 residues with RMSD 4.341 and 3.837 Å, respectively (Fig. 5*B*). Since these RMSD values are high because CE works with all structural regions in the superpositions, we used TM-Align to set a quantitative degree of fold structural similarity through its TM-score defined in a (0–1) scale, with higher values meaning higher fold similarity (56). Superpositions of AT1G56140 with *X. laevis* and *Arabidopsis thaliana* malectins gave TM-scores 0.623 and 0.710, respectively, indicating highly similar folds of AT1G561*nn* MAL domains to animal malectin and to plant malectin-like structures (Fig. 5*B*).

**Figure 5.**
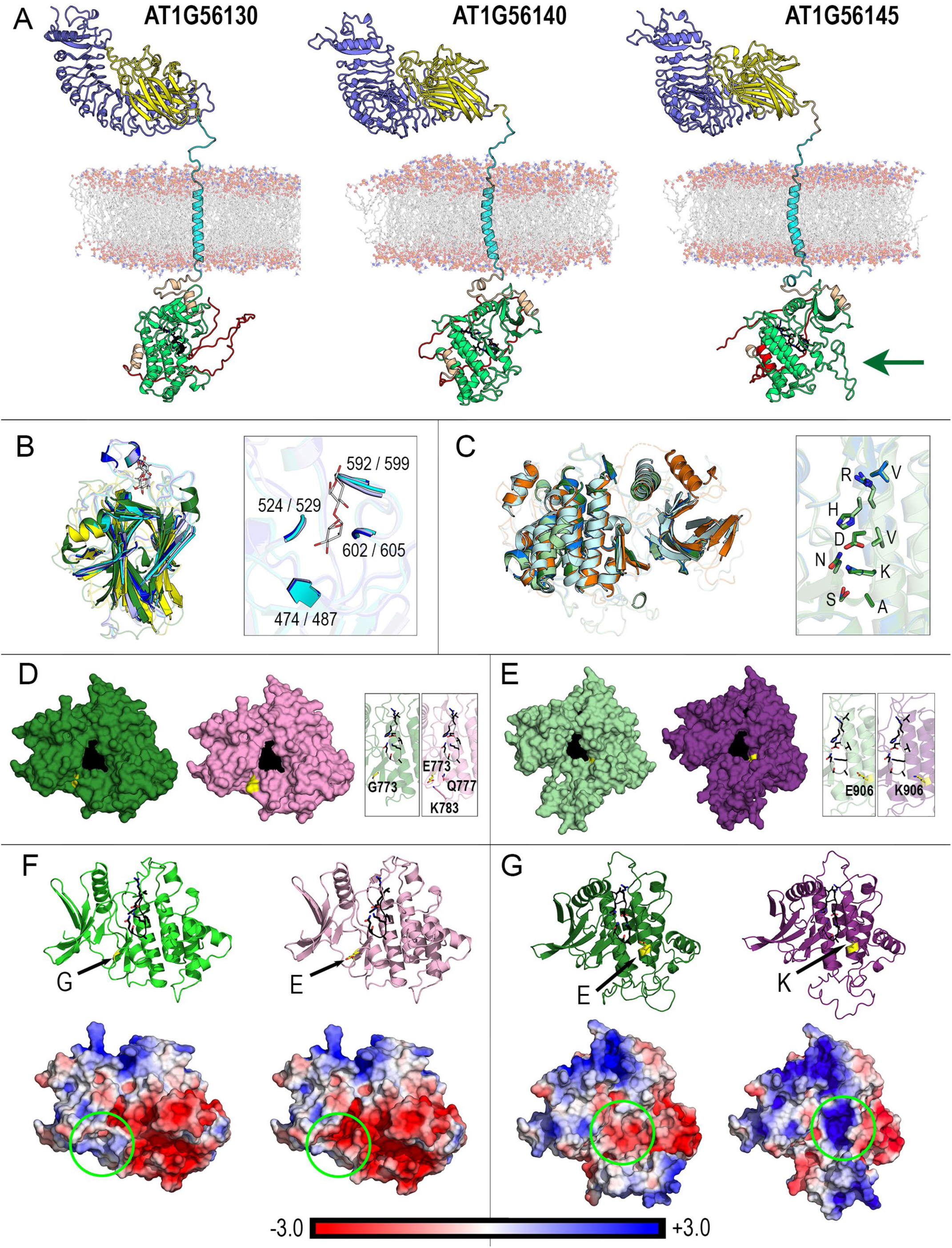
Model structures of AT1G56130 (IGP2/IGP3), AT1G56140 (IGP4) and AT1G56145 (IGP1) LRR-MAL RLKs. (*A*) Structures obtained from AlphaFold models as explained in Material and Methods to suit the extracellular/membrane/intracellular organization. Domains colored as follows: N-terminal, orange; LRR, slate blue; MAL, yellow; segment containing transmembrane helix, cyan; KD, green with catalytic residues shown as black sticks; C-terminal, red. The bilayer modelled by 256 POPC lipids is shown as sticks for carbons and spheres for P, O, and N atoms in polar heads. The arrow in AT1G56145 indicates an extra loop in its KD. (*B*) Left: structural alignment of MAL domains of AT1G56130 (light blue), AT1G56140 (blue) and AT1G56145 (cyan) models with experimental structures of MAL domain from *X. laevis* (yellow; NMR structure PDB code 2KR2) in complex with maltose (sticks with white carbons) and MAL-like domain of ANX1 ECD from *Arabidopsis thaliana* (deep green; crystal structure PDB code 6FIG). Secondary structure elements are shown as opaque cartoons and loops are drawn as partially transparent ribbons to highlight the conserved β-sandwich fold. Right: segments of the three AT1G561*nn* proteins presumably involved in glycan-binding as predicted by selecting residues at 4 Å from maltose in the structural superposition at the left. Labels indicate the sequence positions of those segments in the three proteins. (*C*) Left: Structural alignment of kinase domains of AT1G56130 (deep green), AT1G56140 (marine blue) and AT1G56145 (pale green) models with experimental structures of mitogen-activated protein kinase 3 (MPK3) from *L. donovani* (pale cyan; crystal structure PDB code 4O2Z) and MPK6 from *A. thaliana* (orange; crystal structure PDB code 5CI6). Same depiction of structural elements as in (*B*) to highlight the overall fold of MPK3/6 fold. Right: catalytic site in the three AT1G561*nn* proteins (sticks with carbons in same colors as in the left image) showing the nearly indistinguishable geometry of side chains of conserved residues VHRDVKASN. (*D*) Molecular surface of wild-type (green) and G773E mutation (*igp2/igp3*, pink) structures of KD of AT1G56130. Surface patches of the catalytic site and position 773 are colored black and yellow, respectively. The insets show the catalytic site (black) and residue G773 (yellow) in wild-type (green transparent cartoon) and in mutant (pink transparent cartoon) structures. Q777 and K783 residues within 4 Å from E773 are also shown in the mutant. (*E*). Molecular surface of wild-type (pale green) and E906K mutant (*igp2/igp3*, purple) structures of KD of AT1G56145. Surface patches of the catalytic site and position E906 are colored black and yellow, respectively. The insets show the catalytic site (black) and residue 906 (yellow) in wild-type (pale green transparent cartoon) and in mutant (K906, purple transparent cartoon) structures. (*F*) KD of AT1G56130 comparing the Poisson-Boltzmann (PB) electrostatic potential (EP) mapped onto the protein surface at orientations shown in above cartoons colored green for wild-type and pink for G773E mutant. (*G*) Comparison for KD of AT1G56145 with cartoons colored green for wild-type and purple for E906K mutant. (*F, G*). Arrow indicates the positions of the mutations and the corresponding residue is shown as yellow sticks. Kinase site VHRDVKASN (815-823 in AT1G56130 and 835-843 in AT1G56145) is shown as black sticks. Green circles indicate surface regions at which the mutation provokes a significant electrostatic change. Bar gives the color code for the PB EP in *kT*/*e* units (*k*: Boltzmann constant, *T*: absolute temperature, *e*: electron charge unit).

Similar structural comparison analyses were performed for the KDs of AT1G56130, AT1G56140 and AT1G56145, whose number of residues are 269, 269, and 302, respectively. We narrowed down the search in the PDB to some kinase crystals like MPK6 from Arabidopsis (5CI6 (57) and 6DTL (58)). Their pair CE superpositions included 256 residues in the three cases with RMSD values (Å): AT1G56130/AT1G56140 = 0.315, AT1G56130/AT1G56145 = 0.578 and AT1G56140/AT1G56145 = 0.622. These values indicate that AT1G56145 kinase domain is noticeably different from that of the two other RLKs due to the extra loop seen in the intracellular part of its complete structure (indicated by the arrow in Fig. 5*A*). The local confidence AlphaFold metric reveals a clear difference in AT1G56145 for which *low* or even *very low* predicted *L*ocal Distance Difference Test (p*l*DDT) values appear in the middle of its KD (*SI Appendix*, Fig. S11*A*-*C*), a feature not found in AT1G56130 or AT1G56140 (*SI Appendix*, Fig. *S11A-C*). This notwithstanding, the catalytic site is nearly indistinguishable in the three proteins (Fig. 5*C*). Next, we tested *in silico* the possible structural impact of single mutations G773E in AT1G56130 and E906K in AT1G56145 KDs. Model structures of the KD of wild-type and mutants were newly generated from the corresponding sequence segments with AlphaFold2 and I-TASSER, to test that the backbone in fact remained unaltered though we found differences in loop flexible regions, minor in AT1G56130 and greater in AT1G56145. The conformation of side chains of catalytic residues was identical in wild-type and mutant structures (Fig. 5*D,E*), a result worth emphasizing as the mutation positions are near the catalytic site in both proteins. The changes G773E in AT1G56130 and E906K in AT1G56145 changes had the effect of increasing the surface patch associated with the mutated position (Fig. 5*D,E*). In the case of AT1G56130, the G773E mutation also brought two amino acids (Q777 and K783) into a close vicinity of 4 Å from the mutated residue (Fig. 5*D*) whereas in AT1G56145 the E906K mutation had no significant variations in the neighborhood of the mutated residue (Fig. 5*D*). The major effect of these single mutations was found in the surface electrostatic potential (Fig 5*F,G*). In wild-type AT1G56130, there are 11 residues (7 non-polar, 3 polar, and 1 positively charged, K783) at 5 Å from G773 that confer a weakly positive electrostatic character to a large area around position 773 that becomes strongly negative upon the G773E mutation (Fig. 5*G*). In wild-type AT1G56145 there are 15 residues (12 non-polar, 2 polar, and 1 positively charged, R911) at 5 Å from E906 that produce a weakly negative electrostatic character, which is besides neutralized in part by R911 (Fig. 5*G*). In the mutant, the negative charge is substituted for a positive charge and the new K906 together with R911 give place to a strongly positive electrostatic potential in a large area around position 906 (Fig. 5*G*). Because of these local distributions of residues, the electrostatic effects extend over a surface region far larger than that expected from the small, exposed surface areas of residues 773 (Fig. 5*D*) and 906 (Fig. 5*E*). These changes in KD domains of IGP1/AT1G56130 and IGP2/IGP3/AT1G56130 LRR-MAL RLKs might explain their loss of functionality.

### The ECD of AT1G56145 directly binds cellulose derived oligosaccharides CEL3 and CEL5

Based on the initial structural models of MAL domains of the three LRR-MAL RLKs and their similarities with MAL domain from *Xenopus* sp. that binds short glycans ((53) and Fig. 5*B*), we tested whether ECDs of these LRR-MAL RLKs could be glycan receptors for cellulose and MLG derived oligosaccharides. We expressed the ECDs of AT1G56145 and AT1G56140 in insect cells and purified them by affinity chromatography (*SI Appendix*, Fig. S12*A*). Then, ITC experiments (59) were carried out to test the binding of MLG43 and CEL3 to these ECDs. ITC results proved direct interactions of CEL3 with the ECD of AT1G56145 (Kd = 1.19 ± 0.03 μM; Fig. 6*A*), but not with AT1G56140 (*SI Appendix*, Fig. S12*B*). Results obtained in ITC clearly indicated that these ECDs did not bind, at least directly, to MLG43 (Fig. 6*A, B* and *SI Appendix*, Fig. S12*B*). Similar binding experiments were performed with the ECD of AT1G56145 and CEL5 to determine the specificity of receptor-ligand recognition, and direct binding was also detected with similar high affinity (Kd = 1.40 ± 0.01 μM: Fig. 6*A*). The binding reaction measured for CEL3 and CEL5 were exothermic, with a single binding site (N = 1) and very similar values of ΔH, indicating that extra sugar subunits in the CEL5 oligomer do not improve the detected binding. These data support the role of the ectodomain of AT1G56145 as a receptor for cellulose-derived oligosaccharides and suggest that AT1G56140 RLK might function as a co-PRR in the sensing complex for cellulose and MLG derived oligosaccharides.

**Figure 6.**
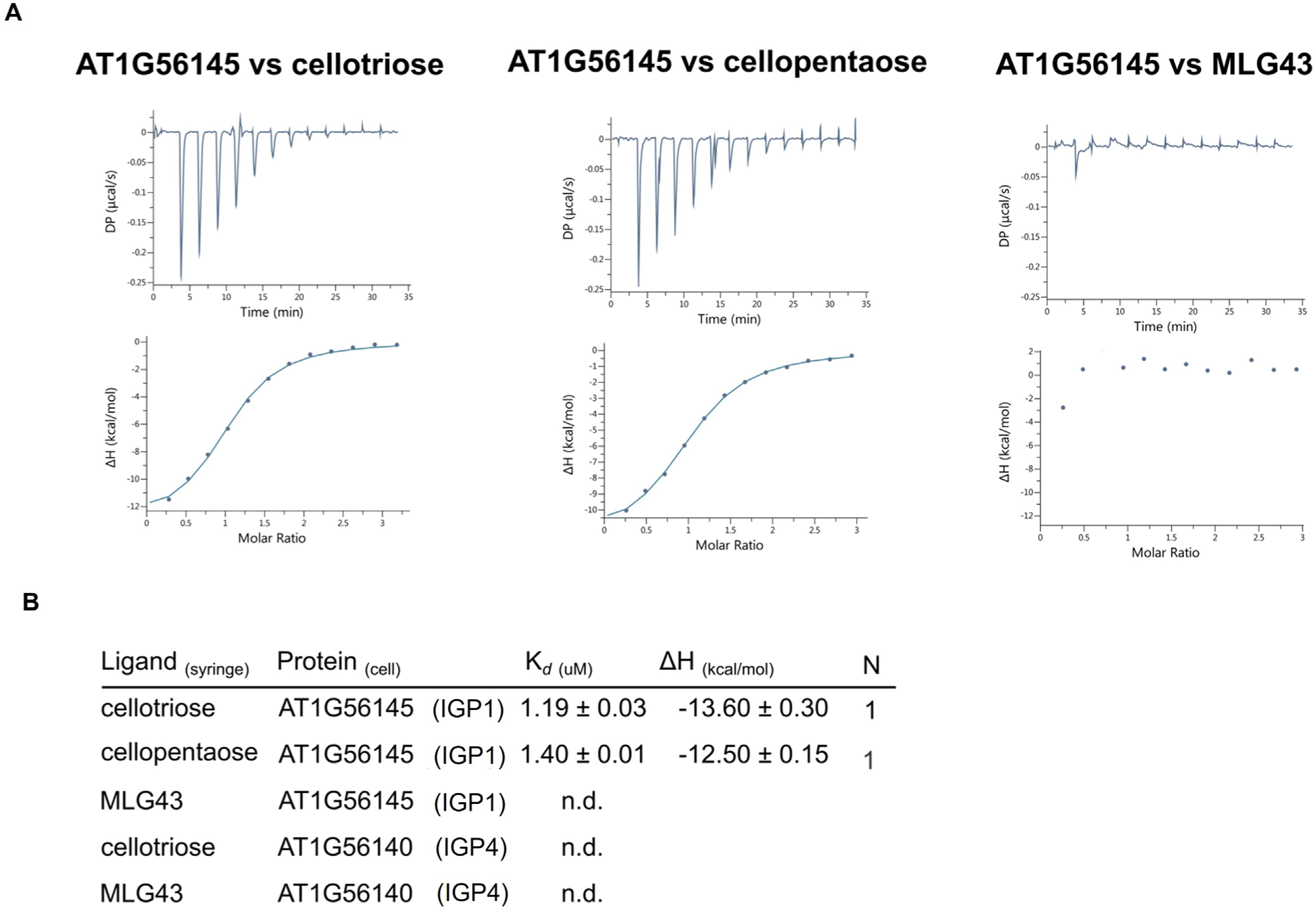
AT1G56145 ECD directly binds CEL3 and CEL5 oligosaccharides. (*A*) Isothermal titration calorimetry (ITC) experiments of AT1G56145 ECD vs. CEL3, MLG43 and CEL5. (*B*) ITC table summaries of AT1G56145 and AT1G56140 vs CEL3, MLG43 and CEL5 oligosaccharides. The binding affinities of AT1G56145 ECD and the corresponding glycans are reported as K*_d_* (dissociation constant, in micromoles), N indicates the reaction stoichiometry (N = 1 for a 1:1 interaction) and ΔH indicates the enthalpy variation. Values indicated in the table are means ± S.D. of independent experiments (n = 2). n.d indicates no binding detected.

## DISCUSSION

Plant immunity is activated by a diverse set of ligands (DAMPs/MAMPs) of different biochemical nature and structure (e.g., peptides and carbohydrates) that are specifically perceived by diverse sets of ECDs from plant PRRs. The specific and direct recognition of a DAMP/MAMP by the ECD of its PRR counterpart receptor trigger the formation of a ligand-PRRs complex, that involves additional PRRs that directly (co-receptors) or indirectly contribute to the functionality of the recognition complex through the stabilization of ligand-receptor binding or activation of KD of some PRRs from the complex, which in turn initiates phosphorylation events and PTI-associated responses (1–3). Several new glycoligands (DAMPs/MAMPs) derived from cell walls or extracellular matrixes from plants or microorganisms that trigger immune responses in plants have been recently described ((9–20), (31–34)). These new carbohydrate-based elicitors include oligosaccharides derived from plant cell wall β-glucans, like MLGs (e.g., MLG43, MLG34 and MLG443) or cellulose (e.g., CEL2, CEL3, CEL4, CEL5, CEL6), or from arabinoxylans and mannans polysaccharides (16, 18, 31–33). These glycoligands have been added to the list of previously characterised oligosaccharides, which include oligoglycans from fungal chitin (e.g., CHI6) or plant pectins (e.g., OGs), which are perceived by LysM-PRRs and WAK1/2/FER1-PRRs, respectively (40, 47, 48). Here, we describe a new group of *Arabidopsis thaliana* RLKs (AT1G56145 (IGP1), AT1G56130 (IGP2/IGP3), and AT1G56140 (IGP4)) with LRR-MAL domains in their ECDs, which function as PRRs triggering immune responses mediated by oligosaccharides derived from cellulose (e.g., CEL3-CEL6) and MLGs (e.g., MLG43 and MLG34; Fig. 1 and Fig. 2). Remarkably, we also demonstrate that the ECD of AT1G56145 (IGP1) directly binds CEL3 and CEL5 with high affinity (Kd = 1.19 ± 0.03 μM, and Kd = 1.40 ± 0.01 μM, respectively), but it does not bind MLG43 (Fig. 6 and S*I Appendix. Fig. S12*). These data suggest that these LRR-MAL RLKs have a function in *Arabidopsis thaliana* as either receptor (AT1G56145 for CEL3/CEL5) or components of the PRR complex involved in the perception of these oligosaccharides or the signaling complex activating PTI responses triggered by the oligosaccharides (Fig. 4). The function of AT1G56145 RLKs as a receptor for cellulose derived oligosaccharides has also been recently proposed by Tseng *et al*., (2022; bioRxiv preprint (60)), that identified the Cellulose Oligosaccharide Receptor Kinase 1 (CORK1) in a similar genetic screening to the one performed here; but without reporting any binding of ECD-CORK1 to cellulose-derived oligosaccharides. We now demonstrate that AT1G56145 RLKs is a true PRR receptor for CEL3/CEL5 oligosaccharides further corroborating its function in *Arabidopsis thaliana* immunity. In a previous independent approach, a Poly(A) ribonuclease (AtPARN, AT1G55870) was found to be also required for the regulation of the immune responses triggered by CEL3 in *Arabidopsis thaliana*, probably through a mechanism of degradation of polyA tail from specific mRNA (61). However, this regulatory mechanism of CEL3-mediated responses acts downstream of the CEL3 perception by PRRs at the membrane surface described here.

Our results support the addition of LRR-MAL RLK family to the set of plant PRRs that are involved in the perception of carbohydrate-based DAMPs/MAMPs, as recently hypothesized (6). This set of PRRs includes members of LysM, WAK and CrRLK1 RLK families (5, 6, 43, 44, 47, 48, 62). The function of the majority of these proteins in oligosaccharides perception and PTI activation has been discovered through the isolation and characterization of *Arabidopsis thaliana* mutants (e.g., *cerk1, lyk4* and *lyk5, wak1* and *wak2, or fer1*), which are generally only partially impaired in PTI responses triggered by specific DAMPs/MAMPs, due to their redundant functions with additional PRRs. Accordingly, PTI deficient phenotypes in these PRRs mutants need to be validated in double or triple mutants (40, 43, 44, 47, 62–64). Similarly, *AT1G56145* (*IGP1*), *AT1G56130* (*IGP2/IGP3*) and *AT1G56140* (*IGP4*) described here were discovered in a screening of *Arabidopsis thaliana* mutants impaired in glycan perception (*igp*), that led to the identification of nine (*igp1-igp9*) mutants, that initially were found to be impaired in MLG43 recognition, and later proved to be also defective in the perception of cellulose-derived oligosaccharides (CEL3-CEL5: Fig. 3 and *SI Appendix Fig S3*). These results suggested that the mechanisms of perception of CEL/MLG oligosaccharides are quite similar in *Arabidopsis thaliana,* and this hypothesis was confirmed by CEL3/MLG43 cross-elicitation experiments (S*I Appendix. Fig. S8*). Despite the similarities between the mechanisms of perception of CEL and MLG-derived oligosaccharides in *Arabidopsis thaliana*, some differences in these mechanisms might exist. For example, we found that LysM-RLKs (i.e., CERK1, LYK4 and LYK5) have a partial contribution in the perception of MLG43 (*SI Appendix Fig S9*), as described previously (31). In contrast, their role in CEL3 perception is minor since PTI activation was observed to be quite similar in *cerk1-2, cer1-2lyk4lyk5* and Col-0 plants treated with CEL3 (S*I Appendix. Fig. S9*). The partial requirement of LysM-RLKs for MLG perception in *Arabidopsis thaliana* is in line with the described function of rice LysM-RLKs members in the perception of MLG oligosaccharides and immune activation. OsCERK1 has been suggested to be the PRR receptor of MLG oligosaccharides based on Microscale Thermophoresis analysis performed with ECD produced in baculovirus, whereas OsCeBiP, the rice receptor for chitin oligosaccharides, has been proposed to be the MLGs co-receptor based on the lack of binding of its ECD to these ligands (33). K*_d_* values of ECD-OsCERK1 for MLG43 are in the 1-2 μM range, similar to that found for AT1G56145 (IGP1) in this work (Fig. 6) and to that of ECD-OsCeBiP for chitohexaose (33). It is of note, that perception of CHI6 in *Arabidopsis thaliana* is not altered in *igp1-igp4* impaired in LRR-MAL RLKs nor in *igp5*-*igp9* mutants (Fig, 1 and *SI Appendix Fig. S2*), further indicating that IGP1-IGP9 proteins are not required for CHI6 perception. These data indicate that at least two different mechanisms for the perception of CHI6 and CEL3/MLG43 oligosaccharides might exist in *Arabidopsis thaliana*. The mechanisms of PTI activation mediated by IGP1, IGP2/IGP3 and IGP4 LRR-MAL RLKs also differ from that of pectin and OGs perception involving FER1 and WAK1-WAK2 (47, 48, 63), respectively, since Ca^2+^ burst in *igp1*^AEQ^*-igp4*^AEQ^ upon OGs treatment was similar to that of Col-0^AEQ^ plants (Fig. 3). Similarly, response to other MAMPs/DAMPs in *igp1*^AEQ^*-igp4*^AEQ^ seedling is not impaired in comparison to that in wild-type plants, further supporting the specific function of IGP1, IGP2/IGP3 and IGP4 RLKs in MLG/CEL oligosaccharide recognition (Fig. 3).

LysM-PRR members (e.g., CERK1/LYK5/LYK4 in *Arabidopsis thaliana* or CeBiP/CERK1 in rice (44, 62) are the only PRRs from plants that have been demonstrated to directly bind oligosaccharides (i.e., CHI4-CHI8 or MLGs) either by ITC or Microscale Thermophoresis analysis performed with purified ECDs or by the generation of crystalized structures ECDs-CERK1/chitin oligosaccharides, which have revealed the structural based of this recognition (33, 43). Other plant ECD-RLKs, like ANX1 and ANX2 that harbour two tandem MAL domains, have been also purified and crystalised, but their putative glycoligand(s) has not been identified (54). Notably, MAL domains like those of ANX1/ANX2 are present in the ECD of the three IGPs RLKs identified in this work, that also contains a LRR domain in the N-terminal (Fig. 5). We have generated *de novo* structural models for the three LRR-MAL RLKs, and these models suggest that AT1G56145 (IGP1), AT1G56130 (IGP2/3) and AT1G56140 (IGP4) have similar structures, in line with their putative recent evolutionary divergence (*SI Appendix Fig. S3A*), and that AT1G56145 has an extra loop in its KD (Fig. 5). Remarkably, MAL domains of these RLKs are structurally very similar to that of *Xenopus* sp. MAL protein, that has been involved in oligosaccharide binding, and to that of plant ANX1/ANX2 from CrRLK1 family (Fig. 5). We hypothesize that the LRR domain of ECDs in IGP1/AT1G56145 might be essential for the formation of the structural pocket involved in CEL3 and CEL5 binding, since glycoligands for ANX1/ANX2 MAL domains have not been identified (65), though this hypothesis deserve experimental confirmation. In the recent article by Tseng *et al.*, (60), two Phe residues conserved in all *Arabidopsis thaliana* MAL domains, but not in MAL from *Xenopus sp.* (53) has been suggested to be essential for CEL3 perception and PTI activation, but this hypothesis has not been validated through binding experiments. The obtention of crystalized structures of CEL3/ECD-AT1G56145 will contribute to decipher the structural bases of CEL3-CEL5 recognition by LRR-MAL ECD, and to determine if it shows some similarities with the mechanisms of cellulose polymer binding by cellulases from microorganisms. Despite our recent progresses in the characterization of the mechanisms of perception of glycoligands by plants PRRs, our current knowledge of innate immunity activation by oligosaccharides in plants is far away of our understanding of these mechanism in mammals (8, 38, 39). Several carbohydrate-recognition domains (CRDs) motifs have been characterised in mammals that binds glycans that are either identical or quite similar to those perceived by plant PRRs (38, 39). However, CEL3 and MLG43 have not been described to be perceived by mammal innate immunity system, and structural comparison of CRDs and LRR-MAL ECDs is not feasible.

Interestingly *igp1* and *igp2/igp3* mutants have point mutations in their KDs that clearly impact on their functionality (Fig. 2). G773E in AT1G56130 and E906K in AT1G56145 do not seem to impair the catalytic sites of these RLKs which are predicted to be almost identical to that of wild-type KDs (Fig. 5*C*). However, these point mutations are predicted to increase the surface patch associated with the mutated position (Fig. 5*D,E*), having a major effect on the surface electrostatic potential of the residues around the mutated positions, that extends over a surface region far larger than that expected from the small areas corresponding to the exposed surfaces of residues (Fig. 5*F,G*). Since kinase activity can be altered by mutations at distant residues from the active site (66), we can hypothesize that G773E and E906K mutations might either affect the catalytic activity of the KDs or the observed changes in the surface patches and electrostatic potential in the mutated KDs might interfere with the interaction of the RLKs KDs with other PRRs or additional proteins that form part of PRR recognition complexes.

Additional protein families with diverse domains in their ECDs have been suggested to putatively function as receptors or co-receptors in the perception of oligosaccharides MAMPs/DAMPs based of their ECD domains (6), but these proteins (RLKs/RLPs/RPe) are poorly characterized. This myriad of potential PRRs encoded by plant genomes would be needed to perceive the diversity of oligosaccharides that can be released from plant and microbial cell walls by the activity of cell wall degrading enzymes secreted by pathogens during colonization or by plant cells during the interaction with pathogen aiming to degrade cell wall from pathogens to compromise their survival (67, 68). The relevance of these carbohydrate-based ligands released during plant-microbe interaction in the regulation of plant disease resistance has been proved in different plant species that show enhanced disease resistance to different pathogens upon exogenous application of glycans DAMPs/MAMPs (e.g., MLG43) in comparison to untreated plants (15, 31). This oligosaccharide-mediated priming effect on PTI and disease resistance has driven the development of sustainable crop protection solutions based on mixtures of active glycans (DAMPs/MAMPs: (69, 70)). The discovery of counterpart receptors for these active glycans in crops will accelerate the selection of the corresponding genes in breeding programs to enhance crops disease resistance. Notably, *AT1G56145 (IGP1), AT1G56130 (IGP2/3)* and *AT1G56140 (IGP4)* genes are in a cluster of *Arabidopsis thaliana* genome, indicating recent duplication events. These genes form part of a family of at least 12 members that has not been characterized in detail previously, except for RFK1 protein that has been involved in pollen tube growth (51). Interestingly, orthologs of *AT1G56145 (IGP1), AT1G56130 (IGP2/3)* and *AT1G56140 (IGP4)* do not seem to be present in grasses, like rice and corn, whereas they have been identified in other plant species (*SI Appendix Fig. S10*). However, MLG43, and in a lower extend CEL3, have been described to be perceived by cereals suggesting that additional receptors to LRR-MAL RLKs (e.g., LysM-RLKs for MLGs in rice) would be involved in these perception mechanisms (32, 33).

MLG- and cellulose-derived oligosaccharides can be also released upon alteration of plant cell wall integrity triggered by other stresses (e.g., salt or drought) or during plant development (e.g., cell wall remodelling: (71, 72)). During these processes plant endogenous enzymes can hydrolase cell wall polysaccharides, as described recently for a maize GH17 licheninase that releases MLG43 and other oligosaccharides from MLGs (73). The characterization of the role of glycan-mediated responses in these additional processes need to be determined to understand their cross-regulation with PTI/disease resistance responses and their impact on plant fitness. Also, regulation of the homeostasis of cell wall derived oligosaccharide needs to be characterised. For example, several plant berberine-bridge enzymes have been identified that control CEL oligosaccharide homeostasis by oxidating the anomeric carbon of CEL3-CEL6 oligosaccharides and thus reducing both their activity as DAMPs, and their preference use as a carbon source by fungi (74). In summary, the results describe here contribute to our understanding of the mechanism of perception of oligosaccharides by the plant immune system and expand the set of families of PRRs and the ECD structures involved in ligand recognition and immune activation in plants.

## Material and Methods

### Biological material and growth conditions

All the *Arabidopsis thaliana* lines used in this study were in Columbia-0 (Col-0) background. *Arabidopsis thaliana* plants used for cytoplasmatic Ca^2+^ measurements were grown in 96-well plates (1 seedling per well) and for MAPKs phosphorylation and gene expression analyses were grown in 24-well plates (∼10 seedlings per well). These seedlings were grown under long-day conditions (14 hours of light) at 19-22°C in a liquid MS1/2 medium. Plants were also grown in soil-vermiculite (3:1) mixture under short-day photoperiod (10 hours of ligh-14h ours dark, 21-20 °C) or long-day photoperiod (14 hours of light-10 hours dark, 19-22°C). *Arabidopsis thaliana* lines used in this work were Col-0^AEQ^ (49, 50), *cerk1-2*^AEQ^ (50), *rbohD* (52) and *igp*^AEQ^ isolated in this work. *cerk1-2lyk4lyk5* triple mutant was generated by crossing *cerk1-2* and *ly4lyk5* (31) double mutant and selecting the triple mutants with described oligonucleotides (*SI Appendix Data Set 3*). The *at1g56140 (igp4) and the at1g56120* T-DNA mutants were obtained from NASC (SALK_005808 and SALK_043782, respectively).

### Carbohydrates used in the experiments

MLG43 (β-1,4-D-(Glc)_2_-β-1,3-D-Glc, # O-BGTRIB), MLG34 (β-1,3-D-(Glc)_2_-β-1,4-D-Glc,; # O-BGTRIA), hexaacetyl-chitohexaose (CHI6; β-1,4-D-(GlcNAc)_6_; #O-CHI6), CEL3 (β-1,4-D-(Glc)_3_; O-CTR), CEL4 (β-1,4-D-(Glc)_4_; # O-CTE), CEL5 (β-1,4-D-(Glc)_5_; # O-CPE) were purchased from Megazyme (Wicklow, Ireland). CEL2 (cellobiose, β-1,4-D-(Glc)_2_; C7252) was purchased from Sigma-Aldrich. OGs (Galacturonan oligosaccharides Mixture DP10/DP15; GalAα1-4[GalAα1-4]8_-13_GalA; GAT114) were purchased from Elicityl (Crolles, France). The peptides Flg22 and AtPEP1 were synthesized from EZBiolab (Carmel, USA) and Abyntek (Zamudio, Spain), respectively.

### Aequorin luminescence measurements

*Arabidopsisthaliana* 8-day-old liquid-grown seedlings carrying the calcium reporter aequorin (*35S::Apoaequo_cyt_* in Col-0 background, Col-0^AEQ^; (31, 49)) were used as described for cytoplasmic calcium (Ca^2+^) measurements in the genetic screening performed to identify the *igp*s and to test the specificity of the *igp*s to different compounds, as described previously (31). Negative controls (water) were included in all the experiments. Aequorin luminescence was recorded with a Varioskan Lux Reader (Thermo Scientific).

### Genetic screening to identify *igp* mutants

*Arabidopsis thaliana* Col-0^AEQ^ seeds were mutagenized with 0,3% EMS for 17 hours. Seeds were sown in soil-vermiculite (3:1) to obtain next-generation seeds (e.g., M1 and M2) (75). M2 seedlings were grown *in vitro* for 8 days, and calcium influxes were evaluated using a luminometer upon treatment with 100 μM MLG43. The bioluminescence of each seedling was analysed individually and those with a low response to the treatment were transferred to soil (230 putative mutants out of 6,400 total seedlings screened), self-crossed and then Ca^2+^ burst tested in F1 seedlings to confirm the impaired response to MLG43. The most interesting (lower response) and validated *igp* mutants were selected for further characterization and backcrossed with Col-0^AEQ^ (*igp1*^AEQ^-*igp3*^AEQ^ and *igp5*^AEQ^*-igp9*^AEQ^ described in this work, and *igp10*^AEQ^*-igp20*^AEQ^, which are not described here). Total calcium discharge was performed by treating seedlings with1M CaCl_2_ and Ca^2+^ burst then was measured in the Luminometer.

### Mapping by whole Genome Sequencing and SNP analysis

Whole-genome sequencing of F2 *igp*^AEQ^ x Col-0^AEQ^ segregating plants with impaired response to MLG43 led to the identification of Single Nucleotide Polymorphism (SNPs) associated with phenotypes in each single mutant isolated. Tissue from 50 F2 segregant individuals was harvested and pooled for gDNA sequencing. gDNA from control line Col-0^AEQ^ was also sequenced as a reference for the SNPs analysis. DNA of the samples were introduced in FastQC v.0.11.9 for quality report (76) and then whole-genome sequencing (150 bp pair-end reads) was performed in an Illumina platform to reach a coverage of 30 million reads (Macrogen, Seoul, South Korea). Reads were aligned with BWA-MEM v.0.7.17 against *Arabidopsis thaliana* TAIR10 genome release (77). BAM files were obtained with samtools v.1.15.1. Variant Caller 16GT Docker image was employed to obtain VCF files (78, 79). From these VCF files, with bcftools v.1.15.1 chromosome, position, reference, alternate and allelic depth (AD) fields were extracted and placed in Excel files. Col-0^AEQ^ SNPs were subtracted from the resulting files to obtain mutant exclusive SNPs. Frequency was calculated from AD fields as follows (AD-alternate allele / (AD-alternate allele + AD-reference allele)) and those SNPs with frequency values higher than 0.99 were selected for further analyses (*SI Appendix Dataset 2*).

### Reactive oxygen species determination

Ten-day-old seedlings of *Arabidopsis thaliana* Col-0 or the mutants were grown *in vitro* conditions as explained above. H_2_O_2_ production was determined after treatments with MAMPs/DAMPs using the luminol assay and a microplate luminescence reader (31).

### Immunoblot analysis of MAPK activation

Twelve-day-old *Arabidopsis thaliana* seedlings grown on liquid MS1/2 medium in 24-well plates were treated with water (mock) and different oligosaccharides for 0, 10 and 20 minutes, and then seedlings were harvested in liquid nitrogen and homogenized using FastPrep Bead Beating Systems (MP Biomedicals) in extraction buffer (50 mM Tris-HCl pH 7.5, 200 mM NaCl, 1 mM EDTA, 10 mM NaF, 2 mM sodium orthovanadate, 1 mM sodium molybdate, 10% (v/v) glycerol, 0.1% (v/v) Tween-20, 1 mM 1,4-dithiothreitol, 1 mM phenylmethylsulfonyl fluoride, and phosphatase inhibitor cocktail P9599 (Sigma)). Total protein extracts were quantified by Bradford assay (Bio-Rad). Protein electrophoresis and Western-Blot were performed as described previously using the Phospho-p44/42 MAPK (Erk1/2) (Thr202/Tyr204) antibody (Cell Signaling Technology) (1:1000) (31). After a membrane stripping process consisting of 5 minutes incubation in a stripping buffer (50 mM glycine pH=2, 0,1% SDS, and 0,05% Tween 20), followed by 10 washing steps and a one-hour blocking in TBS-T with 1% non-fat milk, the same membranes were incubated with Anti-AtMPK3 (1:2500) and horseradish peroxidase-conjugated anti-rabbit antibody (GE-Healthcare) (1:5000). The membranes developed following such protocol was used as a total protein loading control.

### Gene expression analyses

Gene expression analysis was carried out in 12-day-old *Arabidopsis thaliana* seedlings grown on liquid MS1/2 medium after treatment with oligosaccharides (i.e., 100 µM MLG43, 10 µM CEL3, 50 µM CHI6) or water (mock) solutions for 0- and 30-minutes Total RNA was extracted from 100 mg of frozen and ground tissue from mock and treated seedlings using the RNeasy Plant Mini Kit (Qiagen) according to the manufacturer’s protocol (31). qRT-PCR analyses were performed as previously reported (31). *UBC21* (*AT5G25760*) expression was used to normalize the transcript level in each reaction. In the case of showing normalized data to mock conditions, the expression and normalization were determined using the Pfaffl (80). Oligonucleotides used for PCR are described in *SI Appendix Dataset 4*.

### Phylogenetic analysis

The evolutionary history of IGP proteins was inferred using the Minimum Evolution method (81). The bootstrap consensus tree inferred from 1000 replicates is taken to represent the evolutionary history of the taxa analyzed (82). Branches corresponding to partitions reproduced in less than 50% bootstrap replicates are collapsed. The percentage of replicate trees in which the associated taxa clustered together in the bootstrap test (1000 replicates) are shown next to the branches (82). The evolutionary distances were computed using the p-distance method (83) and are in the units of the number of amino acid differences per site. The ME tree was searched using the Close-Neighbor-Interchange (CNI) algorithm (83) at a search level of 1. The Neighbor-joining algorithm (84) was used to generate the initial tree. The analysis involved 25 amino acid sequences. All ambiguous positions were removed for each sequence pair. There were 709 positions in the final dataset. Evolutionary analyses were conducted in MEGA6 (85).

### Structure analyses *in silico*

Model structures of AT1G56130, AT1G56140 and AT1G56145 were downloaded from the AlphaFold Protein Structure Database (86). These models present six identifiable domains: N-terminal containing a signal peptide annotated in Pfam (87), LRR, MAL, TM, KD, and C-terminal tail. The p*l*DDT metricover value of most of LRR, MAL and KDs is ≥ 90% (*SI Appendix*, Fig. S11*A-C*) indicative of a reliability *a priori* compared to experiment (88). To achieve the proper extracellular/TM/intracellular domains separation and taking the AT1G56140 model as benchmark, we proceeded as follows: 1) Torsions were applied to backbone dihedral angles in the segment following MAL with UCSF Chimera 1.15 (89); 2) Energy minimization of the extended segment joining MAL and TM domains was performed with Chimera 1.15 keeping fixed all the remaining structure; 3) The resulting structure was inserted in a pre-equilibrated model of a bilayer composed of 256 POPC lipids with an 8 Å-radius pore downloaded from the CHARMM-GUI Archive (90) and the protein-bilayer system was parametrized using the CHARMM 3.6 force field (91) with CHARMM-GUI (92); 4) The protein-bilayer system was solvated at 16 Å-margins solvation box and NaCl 0.150 M salt ions with VMD 1.9.3 (93), and the structure of the whole protein-bilayer-water-ions system was optimized at 10,000 conjugated gradient energy minimization steps with NAMD 2.14 (94). The optimized final structure of AT1G56140 was then used as input for modeling the corresponding structures of AT1G56130 and AT1G56145 with SWISS-MODEL (95) in “user template mode”. The structural comparisons of MAL and KD in the three AlphaFold models were analyzed with the *Combinatorial Extension* (CE) algorithm (96) implemented in PyMOL 2.5.1 (PyMOL, (97)). To further assess structural fold similarities, the TM-Align web server (56) was also used. The structural analysis of mutants AT1G56130-G773E and AT1G56145-E906K was addressed by modelling separately their KD structures from their sequence segments with both AlphaFold (88) using ColabFold (98) and with I-TASSER (99). Wild-type KDs of both proteins were also modelled with the same methodology in order to double-check the possible structural impact of the mutations.

### Calculation of Poisson-Boltzmann (PB) electrostatic potentials (EPs)

PB EPs were computed with the Adaptive Poisson Boltzmann Solver (APBS) 3.0.0 program (100) through its plug-in in PyMOL 2.5.1 (PyMOL, (97)) using the following settings: nonlinear PB equation solved in sequential focusing multigrid mode, 3D grids of 161^3^ = 4,173,281 points (∼0.5 Å step size), *T* = 298 K, ionic concentration 0.150 M (NaCl), and dielectric constants 4 for proteins and 78.54 for water. The numerical output was saved in OpenDX format and the PB EP was then mapped onto molecular surfaces computed and rendered with PyMOL 2.5.1 (PyMOL, (97)).

### Protein expression and purification from insect cells

Codon-optimized synthetic genes corresponding to the ectodomains of AT1G56145 and AT1G56140 (residues 25 to 630 and 29 to 636, respectively) from Invitrogen GeneArt were cloned into a modified pFastBac donor vector (Geneva Biotech, Geneva, Switzerland) harboring the *Drosophila BiP* (101) or the 30K *Bombyx mori* (102) secretion signal peptides, and with a TEV (tobacco etch virus protease) cleavable C-terminal StrepII-9xHis tag. Baculovirus vectors were generated in DH10MultiBac *E.coli* Cells (Geneva Biotech). Virus amplification was carried out in *Spodoptera frugiperda* Sf9 cells (Geneart, Thermo Fisher Scientific) and was used to infect *Trichoplusia ni* Tnao38 cells (103) for protein expression (virus with a multiplicity of infection (MOI) of 3). The cells were grown 1 day at 28°C and two days at 22°C at 110□rpm. The secreted proteins were subjected to tandem affinity purification, using Ni^2+^ (HisTrap excel, GE Healthcare, equilibrated in 25□mM KPi pH7.8 and 500□mM NaCl) and Strep (Strep-Tactin Superflow high-capacity, (IBA, Germany) equilibrated in 25□mM Tris pH 8.0, 250□mM NaCl, 1□mM EDTA) columns. Affinity tags were removed using His-tagged TEV protease in a 1:50 ratio at 4 °C overnight. Separation of cleavage tags and aggregated proteins was performed using size-exclusion chromatography on a Superdex 200 Increase 10/300 GL column (GE Healthcare) equilibrated in 20□mM citric acid pH 5.0, 150□mM NaCl. Proteins were analysed for purity and structural integrity by SDS-PAGE.

### Analytical size-exclusion (SEC) chromatography

Analytical size-exclusion experiments were performed on a Superdex 200 Increase 10/300 GL column (GE Healthcare) equilibrated in 20□mM citric acid pH 5.0, 150□mM NaCl. 400 μg of protein (aprox. 6 μM) were injected using a loop of 1 mL, sample was eluted with a flow of 0.5 mL/min. UV absorbance at 280 nm was used to monitor the elution of proteins. The peak fractions were analysed by SDS-PAGE followed by Coomassie blue staining.

### Isothermal titration calorimetry (ITC)

Experiments were performed at 25°C using a MicroCal PEAQ-ITC (Malvern Instruments) with a 200□µL standard cell and a 40□μL titration syringe. Proteins were gel-filtrated into the ITC buffer (20□mM sodium citrate pH 5.0, 150□mM NaCl). A typical experiment consisted of injecting 3 μL of potential ligand (CEL3, CEL5 or MLG43) at a concentration range between 135 and 400□μM into the ITC cell containing AT1G56145 or AT1G56140 protein at 9 μM. A total of 13 injections were done at 150□s intervals and 500□rpm stirring speed. Dilution heat was corrected using as a control the thermograph of titration of the ligand into the cell containing only buffer. Experiments were done in duplicates or triplicates, otherwise specified, and data were analysed using the MicroCal PEAQ-ITC Analysis Software provided by the manufacturer. All ITC runs used for data analysis have an N ranging from 0.98 to 1.05. The N values were fitted to 1 in the analysis

## Supporting information

Martin-Dacal_etal_SI_Figures_Datasets

## ACCESION NUMBERS

The genome assembly data of Col-0^AEQ^, *igp1*^AEQ^, *igp2*^AEQ^ *, igp3*^AEQ^ and *igp4* can be retrieved from the NCBI Sequence Read Archive (SRA) under BioProject ID PRJNA864842 and Biosample accessions SAMN30087195, SAMN30087196, SAMN30087197, SAMN30087198 and SAMN30087199, respectively.

Genes described in this work are: *IGP1/AT1G56145; IGP2/AT1G56130; IGP3/AT1G56130; IGP4/AT1G56140; CERK1/AT3G14840; LYK4/AT2G23770; LYK5/AT2G33580; RBOHD/AT5G47910*.

## ACKNOWLEDGEMENTS

We thank Dr. Sonsoles Martín-Santamaria and Elena Gómez-Rubio (Centro de Investigaciones Biológicas Margarita Salas-CSIC, Spain) for their suggestions and advise to understand LRR-MAL structural ECD and validation of *in silico* calculation. We thank Dr. Jose María Jiménez-Gomez and Dr. René Toribio (Centro de Biotecnología y Genómica de Plantas (CBGP), Spain)) for their advice to carry our SNP identification and Western-blot stripping experiments, respectively. This work was supported by grants RTI2018-096975-B-I00 of Spanish Ministry of Science, Innovation and Universities, to AM. This work has been also financially supported by the “Severo Ochoa (SO) Programme for Centres of Excellence in R&D” from the Agencia Estatal de Investigación (AEI) of Spain (grants SEV-2016-0672 (2017–2021) and CEX2020-000999-S (2022–2025) to the CBGP). In the frame of these SO programs H.M. and P.F.-C. were supported with a postdoctoral fellow and the WALLADAPT Mission project was financed. M.M.-D, D.J.B. and D.R. were recipients of PhD Fellows PRE2019-088120 and PRE2019-091276 (SEV-2016-0672) from AEI, and IND2017/BIO-7800 from Madrid Regional Government, respectively. Research in J.S. lab was financially supported by the University of Lausanne, the European Research Council (ERC) grant agreement no. 716358 and the Swiss National Science Foundation grant no. 310030_204526.

## SUPPORTING INFORMATION

Additional Supporting Information may be found in the online version of this article

## Appendix SI Information

**Supplemental Figure 1. Forward genetic screen using Col-0^AEQ^ was used as tool to discover new PRRs and proteins involved in oligosaccharide perception and PTI activation.**

(*A*) *Arabidopsis thaliana* seeds carrying the *Apoaequorin* transgene (*35S::Apoaequo_cyt_,* Col-0^AEQ^), encoding a calcium cytoplasmatic sensor, were mutagenized with 0,3% (v/v) EMS for 17 hours. T2 seedlings were grown *in vitro* for 8 days, and calcium influxes were evaluated using a luminometer upon oligosaccharide treatment. Seedlings with a low response to the treatments were transferred to soil and backcrossed with Col-0^AEQ^. Whole-genome sequencing of F2 segregating plant for this backcross led to the identification of SNPs associated with the phenotype of each single mutant isolated (*SI Appendix Data Set 2*). (*B*) Total calcium of the *igp1-igp3*^AEQ^ was discharged by the addition of 1mM CaCl_2_ to the wells with the seedlings. The kinetic areas after CaCl_2_ treatment were integrated and their values used for the calculation of the total Ca^2+^ % produced which was similar in all genotypes tested. (*C*) Elevations of cytoplasmatic calcium concentration over time, measured as relative luminescence units (RLU), after treatment with 100 µM of MLG43 of eight-day old seedlings of Col-0^AEQ^ and F1 *igp2*^AEQ^*igp3*^AEQ^ cross. Lack of response to MLG43 in F1 seedlings of *igp2*^AEQ^*igp3*^AEQ^ cross indicates that both mutations are allelic. Data represent the mean ± standard error (n=6 in Col-0^AEQ^ and n=12 in F1 of *igp2*^AEQ^*igp3*^AEQ^). (*B.C*) Data from one of the three experiments performed that gave similar results.

**Supplemental Figure 2. Identification of additional *igp* (*igp5-igp9*) mutants impaired in MLG43 perception.**

Calcium burst was measured as relative luminescence units (RLU) over time in eight-day-old seedlings of Col-0^AEQ^ in comparison with *igp5*^AEQ^*-igp9*^AEQ^ mutants after treatment with (*A*) 100 µM of MLG43 or (*B*) 100 µM of CHI6. Data represent the mean ± standard error (n=3 in the case of Col-0^AEQ^ and n=12 in the case of *igp*^AEQ^ mutants).

**Supplemental Figure 3. LRR-MAL RLK family in *Arabidopsis thaliana* includes additional members, such as IGP4/AT1G56140, that is also required for MLG43 perception.**

(A) Phylogenetic tree of the LRR-MAL RLKs in *Arabidopsis thaliana*. The full-length sequence of *Arabidopsis thaliana* proteins predicted to contain a MAL domain in their ECDs (6) were aligned using the ClustalW algorithm (see Material and Methods). The evolutionary history was inferred using the Minimum Evolution method. The analysis involved 25 protein sequences. All ambiguous positions were removed for each sequence pair. Evolutionary analyses were conducted in MEGA6. (B) Endogenous expression of the four genes (*AT1G56120, AT1G56130 (IGP2/3), AT1G56140, AT1G56145 (IGP1))* of the *IGP* clade family members in Col-0 wild-type plants determined by RT-qPCR in seedling treated (30 minutes) with water (mock), MLG43 (100 µM), CEL3 (10 µM) or CHI6 (50 µM). Gene expression values are relative to the housekeeping gene *UBC21* (*AT5G25769*), and to the expression levels in mock-treated. (*C*) Quantitative RT-PCR analysis of the immunity marker genes *CYP81F2* and *WRKY53* in T-DNA lines of LRR-MAL RLKs members of the same clade (*at1g56120* and *at1g56140*/*igp4)* and Col-0 seedlings after treatment with MLG43 (100 µM). Gene expression values are relative to the housekeeping gene *UBC21* (*AT5G25769*) and are normalized to Col-0 (value of 1). Data represent the mean ± standard error of three technical replicates out of three independent biological replicates (n=3). Statistically significant differences were calculated according to Student’s t-test (\**P* <0.05, ** 0.01< P <0.001).

**Supplemental Figure 4. Developmental phenotypes of *igp* plants and LysM-PRR mutants.**

(*A*) Mature siliques and pedicels from 50-day-old plants of the indicated genotypes. (*B,C*) Rosettes in 23 day-old plants grown under short-day (*B*) photoperiod (10:14 hours light: dark). and 28 day-old plants grown long-day (*C*) photoperiod (14:10 hours light:dark). *c1/l4/l5* is the abbreviation of the triple mutant *cerk1-2lyk4lyk5* generated in this work.

**Supplemental Figure 5. Cytoplasmic calcium burst in response to CEL3 is impaired in *igp5-igp9* mutants.**

Ca^2+^ influx measured as relative luminescence units (RLU) over time in 8-day-old *Arabidopsis* Col-0^AEQ^ and *igp5*^AEQ^*-igp9*^AEQ^ seedlings after treatment with 10 µM CEL3. Data represent the mean ± standard error (n=3 in the case of Col-0 ^AEQ^ and n=12 in the case of *igp*^AEQ^). Data are from one of the three experiments performed that gave similar results.

**Supplemental Figure 6. Calcium burst in response to MLG34 and cellulose oligosaccharides is impaired in igp mutants.**

Ca^2+^ burst measured as relative luminescence units (RLU) over the time in 8-day-old Col-0^AEQ^, *igp1*^AEQ^, *igp3*^AEQ^ and *igp4*^AEQ^ seedlings after treatment with (*A*) 100 µM MLG34, (*B*) 10 µM CEL4, (C) 10 µM CEL5 and (*D*) 10 µM CEL2. Data represent the mean ± standard error (n=3 in the case of Col-0 ^AEQ^ and n=12 in the case of *igp*^AEQ^). Data are from one of the three experiments performed that gave similar results.

**Supplemental Figure 7. Cross-elicitation during the refractory period of calcium burst triggered by MLG43 or CEL3.**

Data show cytoplasmic calcium burst measured as relative luminescence units (RLU) over time in 8-day-old Col-0^AEQ^ seedlings after sequentially treatments with (*A*) 50µM MLG43 and 50µM MLG43, (*B*) 10µM CEL3 and 10µM CEL3, and (*C*) 50µM MLG43 and 10µM CEL3 (blue) and 10 µM CEL3 and 50 µM MLG43 (red). Arrow indicates the application time of the second elicitor within the refractory period of the first elicitation. Data represent the average RLU values of 4 seedlings (n=4) ± standard deviation. Data are from one of the three experiments performed that gave similar results.

**Supplemental Figure 8. ROS production triggered by flg22 and CHI6 in *igp1-igp4* mutants.** Luminol-assay in 10-day-old seedlings of *igp1*^AEQ^*-igp4*^AEQ^ to monitor H_2_O_2_ production over a period of 50 minutes, measured as relative luminescence units (RLU). Col-0 and *rbohD* genotypes were included as controls. MAMPs/DAMPs were added 5 minutes after incubation of the seedling in plates in the Luminometer: (*A*) 1 μM flg22 and (*B*) 100 μM CHI6. Data represent mean ± standard error (n=24). Student t-test showed statistically significant differences (*P* < 0.001) only in the comparison of Col-0 and *rbohD* in both treatments Data are from one of the three experiments performed that gave similar results.

**Supplemental Figure 9. Activation of PTI hallmarks by MLG43, CEL3 and CHI6 in LysM-PRR mutants.**

(*A-C*) Reactive Oxygen Species (ROS) production was monitored as H_2_O_2_ production over a period of 50 minutes in luminol-assays, measured as relative luminiscence units (RLU), in seedlings of the indicated genotypes. Col-0 and *rbohD* were included as controls. Oligosaccharides were added 5 minutes after incubation of the seedling in plates in the Luminometer: (*A*) 100 μM MLG43; (*B*) 10 μM CEL3; (*C*) 100 μM CHI6. Comparison to Col-0 assessed by Student t-test at the time of the Col-0 peak shows statistically significant differences for *cerk1-2lyk4lyk5* and *cerk1-2* (*P* < 0.001) only in response to CHI6 (*C*), whereas *rbohD* showed statistically significant differences (*P* < 0.001) in all treatments. Data represent mean ± standard error (n=24). Data are from one of the three experiments performed that gave similar results. (*D*) Mitogen-activated protein kinases (MAPK) phosphorylation was analyzed in 12-day-old seedlings of Col-0, *cerk1-2* and *cerk1-2lyk4lyk5 (c1l4l5)* treated with either 100 µM MLG43, 10 µM CEL3, 50 µM CHI6 or water (mock). Western-Blot using anti-pTEpY antibody (anti p42/44) for phosphorylated MAPK moieties of seedling samples harvested at different time points (0, 10 and 20 minutes) was performed. Black arrows indicate the position of phosphorylated MPK6 (top), MPK3 (middle) and MPK4/11 (bottom). Anti-MPK3 was used as total protein control to show the loading of each gel. These results are from one representative experiment out of the two performed that gave similar results. (*E,F*) Quantitative RT-PCR analysis in 12-days-old seedlings of Col-0, *cerk1-2* and *cerk1-2lyk4lyk5 (c1l4l5)* genotypes. Expression levels of immune marker genes (*E*) *CYP81F2* and (*F*) *WRKY53* relative to housekeeping gene *UBC21* (*AT5G25769*) at 30 min after application of oligosaccharide100 µM MLG43, 10 µM CEL3, 50 µM CHI6, or water (mock) are shown. Data represent mean ± standard error of three technical replicates out of three independent biological replicates (n=3). Statistically significant differences between MLG43, CEL3 or CHI6 treated *cerk1-2* and *cerk1-2lyk4lyk5 (c1l4l5) versus* treated Col-0 seedlings according to Student’s t-test (* *P* <0.05, ** 0.01 < *P* > 0.001, *** *P* < 0.001).

**Supplemental Figure 10. Phylogenetic analysis of LRR-MAL RLKs in Arabidopsis and other plant species.**

Phylogenetic tree of the LRR-MAL RLKs family members in selected plant species. The LRR-MAL ECD domain of IGP1/AT1G56145 was blasted against Uniprot protein database to retrieve a total of 248 protein ECD sequences that are evolutionary related to AT1G56145. A representative subset of 25 sequences in different plant species was selected to build up this phylogenetic tree. In addition, the outgroup protein ANX1, a member of a distinct clade of plant malectin RLKs, and the *X. laevis* malectin protein Q6INX3 were included in the tree for comparison. The evolutionary history was inferred using the Minimum Evolution method. Evolutionary analyses were conducted in MEGA6 (see Material and Methods).

**Supplemental Figure 11. AlphaFold predictions for AT1G56130, AT1G56140 and AT1G56145 proteins.**

(*A*) Plot of p*l*DDT (predicted *l*ocal Distance Difference Test) for AT1G56130. AlphaFold uses this metric to give a per-residue confidence score between 0 and 100. Horizontal lines and the corresponding colors follow the AlphaFold guidelines to consider the model confidence *very high* (p*l*DDT > 90, blue), *high* (90 > p*l*DDT > 70, light blue), *low* (70 > p*l*DDT > 50, light orange), and *very low* (p*l*DDT < 50, deep orange). According to AlphaFold, regions with p*l*DDT < 50 may be unstructured in isolation. Vertical lines separate the domains. Regions shaded grey correspond to short segments not assigned to any domain. (*B*) Same plot for AT1G56140. (*C*) Same plot for AT1G56145. (*D*) AlphaFold structure predicted for AT1G56140 protein. The domains are correctly predicted but their spatial organization is not consistent with the extracellular (LRR + MAL) – transmembrane (TM) – intracellular (KD) separation. The short loop region enclosed in the magenta box is the segment used to modify the structure for obtaining the proper spatial organization as explained in Methods.

**Supplemental Figure 12. Purification of ECDs of AT1G56140 and AT1G56145 and ITC of AT1G56140 binding of MLG43 and CEL3.**

(A) Analytical size exclusion chromatography (SEC) and SDS-PAGE of the corresponding proteins purified by SEC and used in the Isothermal titration calorimetry (ITC) binding assays. (*B*) ITC experiments of AT1G56140 ECD vs. CEL3 and MLG43.

## REFERENCES

1. B. P. M. Ngou, P. Ding, J. D. G. Jones, Thirty years of resistance: Zig-zag through the plant immune system. The Plant Cell 34, 1447–1478 (2022).

2. F. Boutrot, C. Zipfel, Function, Discovery, and Exploitation of Plant Pattern Recognition Receptors for Broad-Spectrum Disease Resistance. Annual Review of Phytopathology 55, 257–286 (2017).

3. J. Bigeard, J. Colcombet, H. Hirt, Signaling mechanisms in pattern-triggered immunity (PTI). Molecular Plant 8, 521–539 (2015).

4. D. Tang, G. Wang, Receptor Kinases in Plant-Pathogen Interactions: More Than Pattern Recognition. The Plant Cell 29, 618–637 (2017).

5. K. Bellande, J. J. Bono, B. Savelli, E. Jamet, H. Canut, Plant Lectins and Lectin Receptor-Like Kinases: How Do They Sense the Outside? International journal of molecular sciences 18 (2017).

6. I. del Hierro, H. Mélida, C. Broyart, J. Santiago, A. Molina, Computational prediction method to decipher receptor–glycoligand interactions in plant immunity. The Plant Journal 105, 1710–1726 (2021).

7. C. Franck, J. Westermann, A. Boisson-Dernier, Plant Malectin-Like Receptor Kinases: From Cell Wall Integrity to Immunity and Beyond. Annual Review of Plant Biology 69 (2018).

8. L. Bacete, H. Mélida, E. Miedes, A. Molina, Plant cell wall-mediated immunity: cell wall changes trigger disease resistance responses. The Plant Journal 93, 614–636 (2018).

9. O. Klarzynski et al., Linear beta-1,3 glucans are elicitors of defense responses in tobacco. Plant Physiology 124, 1027–1038 (2000).

10. A. Aziz et al., Elicitor and resistance-inducing activities of beta-1,4 cellodextrins in grapevine, comparison with beta-1,3 glucans and alpha-1,4 oligogalacturonides. Journal of Experimental Botany 58, 1463–1472 (2007).

11. H. Kaku et al., Plant cells recognize chitin fragments for defense signaling through a plasma membrane receptor. Proceedings of the National Academy of Sciences of the United States of America 103, 11086–11091 (2006).

12. A. A. Gust et al., Bacteria-derived peptidoglycans constitute pathogen-associated molecular patterns triggering innate immunity in Arabidopsis. The Journal of Biological Chemistry 282, 32338–32348 (2007).

13. C. Denoux et al., Activation of defense response pathways by OGs and Flg22 elicitors in Arabidopsis seedlings. Molecular Plant 1, 423–445 (2008).

14. J. Claverie et al., The Cell Wall-Derived Xyloglucan Is a New DAMP Triggering Plant Immunity in Vitis vinifera and Arabidopsis thaliana. Frontiers in Plant Science 9 (2018).

15. H. Mélida et al., Non-branched β-1,3-glucan oligosaccharides trigger immune responses in Arabidopsis. The Plant Journal 93, 34–49 (2018).

16. H. Mélida et al., Arabinoxylan-Oligosaccharides Act as Damage Associated Molecular Patterns in Plants Regulating Disease Resistance. Frontiers in Plant Science 11, 1210 (2020).

17. A. Voxeur et al., Oligogalacturonide production upon Arabidopsis thaliana-Botrytis cinerea interaction. Proceedings of the National Academy of Sciences of the United States of America 116, 19743–19752 (2019).

18. H. Zang et al., Mannan oligosaccharides trigger multiple defence responses in rice and tobacco as a novel danger-associated molecular pattern. Molecular Plant Pathology 20, 1067–1079 (2019).

19. A. Wanke et al., Plant species-specific recognition of long and short β-1,3-linked glucans is mediated by different receptor systems. The Plant Journal. 102, 1142–1156 (2020).

20. M. Versluys, E. Toksoy Öner, W. Van den Ende, Fructan oligosaccharide priming alters apoplastic sugar dynamics and improves resistance against Botrytis cinerea in chicory. Journal of Experimental Botany 73, 4214–4235 (2022).

21. R. A. Burton, G. B. Fincher, (1,3;1,4)-beta-D-glucans in cell walls of the poaceae, lower plants, and fungi: a tale of two linkages. Molecular Plant 2, 873–882 (2009).

22. J. L. Morgan, J. Strumillo, J. Zimmer, Crystallographic snapshot of cellulose synthesis and membrane translocation. Nature 493, 181–186 (2013).

23. B. Kloareg, Y. Badis, J. M. Cock, G. Michel, Role and Evolution of the Extracellular Matrix in the Acquisition of Complex Multicellularity in Eukaryotes: A Macroalgal Perspective. Genes 12 (2021).

24. S. C. Fry, B. Nesselrode, J. G. Miller, B. R. Mewburn, Mixed-linkage (1-->3,1-->4)-beta-D-glucan is a major hemicellulose of Equisetum (horsetail) cell walls. New Phytologist 179, 104–115 (2008).

25. I. Sørensen et al., Mixed-linkage (1-->3),(1-->4)-beta-D-glucan is not unique to the Poales and is an abundant component of Equisetum arvense cell walls. The Plant Journal : for cell and molecular biology 54, 510–521 (2008).

26. Z. A. Popper, S. C. Fry, Primary cell wall composition of bryophytes and charophytes. Annals of Botany 91, 1–12 (2003).

27. A. A. Salmeán et al., Insoluble (1L→L3), (1L→L4)-β-D-glucan is a component of cell walls in brown algae (Phaeophyceae) and is masked by alginates in tissues. Scientific Reports 7, 2880 (2017).

28. D. Pérez-Mendoza et al., Novel mixed-linkage β-glucan activated by c-di-GMP in Sinorhizobium meliloti. Proceedings of the National Academy of Sciences of the United States of America 112, 757–765 (2015).

29. T. Fontaine et al., Molecular organization of the alkali-insoluble fraction of Aspergillus fumigatus cell wall. The Journal of Biological Chemistry 275, 27594–27607 (2000).

30. F. Pettolino et al., Hyphal cell walls from the plant pathogen Rhynchosporium secalis contain (1,3/1,6)-beta-D-glucans, galacto- and rhamnomannans, (1,3;1,4)-beta-D-glucans and chitin. The FEBS journal 276, 3698–3709 (2009).

31. D. Rebaque et al., Cell wall-derived mixed-linked β-1,3/1,4-glucans trigger immune responses and disease resistance in plants. The Plant Journal 106, 601–615 (2021).

32. S. Barghahn et al., Mixed Linkage β-1,3/1,4-Glucan Oligosaccharides Induce Defense Responses in Hordeum vulgare and Arabidopsis thaliana. Frontiers in Plant Science 12 (2021).

33. C. Yang et al., Poaceae-specific cell wall-derived oligosaccharides activate plant immunity via OsCERK1 during Magnaporthe oryzae infection in rice. Nature Communications 12, 2178 (2021).

34. C. Yang et al., Addendum: Poaceae-specific cell wall-derived oligosaccharides activate plant immunity via OsCERK1 during Magnaporthe oryzae infection in rice. Nature Communications 12, 3945 (2021).

35. C. A. Souza, S. Li, A. Z. Lin, Cellulose-Derived Oligomers Act as Damage-Associated Molecular Patterns and Trigger Defense-Like Responses. Plant Physiol. 173, 2383–2398 (2017).

36. F. Locci et al., An Arabidopsis berberine bridge enzyme-like protein specifically oxidizes cellulose oligomers and plays a role in immunity. The Plant Journal : for cell and molecular biology 98, 540–554 (2019).

37. F. M. Gámez-Arjona et al., Impairment of the cellulose degradation machinery enhances Fusarium oxysporum virulence but limits its reproductive fitness. Science Advances 8, eabl9734 (2022).

38. M. Taylor et al., “Discovery and Classification of Glycan-Binding Proteins ” in Discovery and Classification of Glycan-Binding Proteins C. R. In Varki A, Esko JD, et al., editors, Ed. (Cold Spring Harbor Laboratory Press Cold Spring Harbor (NY), 2022).

39. R. D. Cummings, Schnaar, R.L., Esko, J.D., Woods, R.J., Drickamer, K., Taylor, M.E., Principles of Glycan Recognition (ed. In Varki A, Cummings RD, Esko JD, et al., editors., 2022).

40. A. Miya et al., CERK1, a LysM receptor kinase, is essential for chitin elicitor signaling in Arabidopsis. Proceedings of the National Academy of Sciences of the United States of America 104, 19613–19618 (2007).

41. R. Willmann et al., Arabidopsis lysin-motif proteins LYM1 LYM3 CERK1 mediate bacterial peptidoglycan sensing and immunity to bacterial infection. Proceedings of the National Academy of Sciences of the United States of America 108, 19824–19829 (2011).

42. Y. Desaki et al., OsCERK1 plays a crucial role in the lipopolysaccharide-induced immune response of rice. New Phytologist 217, 1042–1049 (2018).

43. T. Liu et al., Chitin-induced dimerization activates a plant immune receptor. Science (New York, N.Y.) 336, 1160–1164 (2012).

44. Y. Cao, Y. Liang, K. Tanaka, The kinase LYK5 is a major chitin receptor in Arabidopsis and forms a chitin-induced complex with related kinase CERK1. eLife 3 (2014).

45. T. Shimizu et al., Two LysM receptor molecules, CEBiP and OsCERK1, cooperatively regulate chitin elicitor signaling in rice. The Plant Journal : for cell and molecular biology 64, 204–214 (2010).

46. C. Yang, E. Wang, J. Liu, CERK1, more than a co-receptor in plant–microbe interactions. New Phytologist 234, 1606–1613 (2022).

47. A. Brutus, F. Sicilia, A. Macone, F. Cervone, G. De Lorenzo, A domain swap approach reveals a role of the plant wall-associated kinase 1 (WAK1) as a receptor of oligogalacturonides. Proceedings of the National Academy of Sciences 107, 9452–9457 (2010).

48. W. Tang et al., Mechano-transduction via the pectin-FERONIA complex activates ROP6 GTPase signaling in Arabidopsis pavement cell morphogenesis. Current Biology 32, 508–517.e503 (2022).

49. M. R. Knight, A. K. Campbell, S. M. Smith, A. J. Trewavas, Transgenic plant aequorin reports the effects of touch and cold-shock and elicitors on cytoplasmic calcium. Nature 352, 524–526 (1991).

50. S. Ranf, L. Eschen-Lippold, P. Pecher, J. Lee, D. Scheel, Interplay between calcium signalling and early signalling elements during defence responses to microbe- or damage-associated molecular patterns. The Plant Journal 68, 100–113 (2011).

51. H. K. Lee, D. R. Goring, Two subgroups of receptor-like kinases promote early compatible pollen responses in the Arabidopsis thaliana pistil. Journal of Experimental Botany 72, 1198–1211 (2021).

52. J. Morales, Y. Kadota, C. Zipfel, A. Molina, M. A. Torres, The Arabidopsis NADPH oxidases RbohD and RbohF display differential expression patterns and contributions during plant immunity. Journal of Experimental Botany 67, 1663–1676 (2016).

53. T. Schallus, K. Fehér, U. Sternberg, V. Rybin, C. Muhle-Goll, Analysis of the specific interactions between the lectin domain of malectin and diglucosides. Glycobiology 20, 1010–1020 (2010).

54. S. Moussu, S. Augustin, A. O. Roman, C. Broyart, J. Santiago, Crystal structures of two tandem malectin-like receptor kinases involved in plant reproduction. Acta crystallographica. Section D, Structural biology 74, 671–680 (2018).

55. T. Schallus et al., Malectin: a novel carbohydrate-binding protein of the endoplasmic reticulum and a candidate player in the early steps of protein N-glycosylation. Molecular Biology of the Cell 19, 3404–3414 (2008).

56. Y. Zhang, J. Skolnick, TM-align: a protein structure alignment algorithm based on the TM-score. Nucleic Acids Research 33, 2302–2309 (2005).

57. B. Wang et al., Analysis of crystal structure of Arabidopsis MPK6 and generation of its mutants with higher activity. Scientific Reports 6, 25646 (2016).

58. A. Putarjunan, J. Ruble, A. Srivastava, Bipartite anchoring of SCREAM enforces stomatal initiation by coupling MAP kinases to SPEECHLESS. Nat. Plants 5, 742–754 (2019).

59. P. J. Sandoval, J. Santiago, In Vitro Analytical Approaches to Study Plant Ligand-Receptor Interactions. Plant Physiology 182, 1697–1712 (2020).

60. Y.-H. Tseng et al., CORK1, a LRR-Malectin Receptor Kinase for Cellooligomer Perception in Arabidopsis thaliana. bioRxiv 10.1101/2022.04.29.490029, 2022.2004.2029.490029 (2022).

61. J. M. Johnson et al., A Poly(A) Ribonuclease Controls the Cellotriose-Based Interaction between Piriformospora indica and Its Host Arabidopsis. Plant Physiology 176, 2496–2514 (2018).

62. S. Liu et al., Molecular Mechanism for Fungal Cell Wall Recognition by Rice Chitin Receptor OsCEBiP. Structure 24, 1192–1200 (2016).

63. K. Dünser et al., Extracellular matrix sensing by FERONIA and Leucine-Rich Repeat Extensins controls vacuolar expansion during cellular elongation in Arabidopsis thaliana. The EMBO journal 38 (2019).

64. H. Guo et al., FERONIA Receptor Kinase Contributes to Plant Immunity by Suppressing Jasmonic Acid Signaling in Arabidopsis thaliana. Current Biology 28, 3316–3324.e3316 (2018).

65. S. Moussu et al., Structural basis for recognition of RALF peptides by LRX proteins during pollen tube growth. Proceedings of the National Academy of Sciences 117, 7494–7503 (2020).

66. C. L. McClendon, A. P. Kornev, M. K. Gilson, S. S. Taylor, Dynamic architecture of a protein kinase. Proceedings of the National Academy of Sciences of the United States of America 111, E4623–4631 (2014).

67. J. Wan et al., Cell wall associated immunity in plants. Stress Biology 1, 3 (2021).

68. D. Pontiggia, M. Benedetti, S. Costantini, G. De Lorenzo, F. Cervone, Dampening the DAMPs: How Plants Maintain the Homeostasis of Cell Wall Molecular Patterns and Avoid Hyper-Immunity. Frontiers in Plant Science 11 (2020).

69. C. Chaliha, R. A. Field, E. Kalita, “Glycans as Plant Defense Priming Agents Against Filamentous Pathogens” in Plant Defence: Biological Control, J.-M. Mérillon, K. G. Ramawat, Eds. (Springer International Publishing, Cham, 2020), 10.1007/978-3-030-51034-3_4, pp. 99–118.

70. P. Lemke, L. Jünemann, B. M. Moerschbacher, Synergistic Antimicrobial Activities of Chitosan Mixtures and Chitosan-Copper Combinations. International journal of molecular sciences 23, 3345 (2022).

71. . L. Bacete et al., THESEUS1 modulates cell wall stiffness and abscisic acid production in Arabidopsis thaliana. Proceedings of the National Academy of Sciences of the United States of America 119 (2022).

72. N. Gigli-Bisceglia, E. van Zelm, W. Huo, J. Lamers, C. Testerink, Arabidopsis root responses to salinity depend on pectin modification and cell wall sensing. Development (Cambridge, England) 149 (2022).

73. F. J. Kraemer et al., A mixed-linkage (1,3;1,4)-β-D-glucan specific hydrolase mediates dark-triggered degradation of this plant cell wall polysaccharide. Plant Physiology 185, 1559–1573 (2021).

74. M. Benedetti et al., Four Arabidopsis berberine bridge enzyme-like proteins are specific oxidases that inactivate the elicitor-active oligogalacturonides. The Plant Journal : for cell and molecular biology 94, 260–273 (2018).

75. S. Ranf et al., Defense-related calcium signaling mutants uncovered via a quantitative high-throughput screen in Arabidopsis thaliana. Molecular plant 5, 115–130 (2012).

76. S. Andrews (2010) FastQC: A Quality Control Tool for High Throughput Sequence Data.

77. H. Li (2013) Aligning sequence reads, clone sequences and assembly contigs with BWA-MEM. arXiv:1303.3997v2.

78. R. Luo, M. Schatz, S. & Salzberg, 16GT: a fast and sensitive variant caller using a 16-genotype probabilistic model. GigaScience 6(**7**) 1–4 (2017).

79. P. Danecek et al., Twelve years of SAMtools and BCFtools. GigaScience 10 (2021).

80. M. W. Pfaffl, A new mathematical model for relative quantification in real-time RT-PCR. Nucleic Acids Research 29, e45 (2001).

81. A. Rzhetsky, M. Nei, A Simple Method for Estimating and Testing Minimum-Evolution Trees. Molecular biology and evolution 9, 945–945 (1992).

82. J. Felsenstein, CONFIDENCE LIMITS ON PHYLOGENIES: AN APPROACH USING THE BOOTSTRAP. Evolution; international journal of organic evolution 39, 783–791 (1985).

83. M. Nei, S. Kumar, Molecular evolution and phylogenetics (Oxford University Press, USA, 2000).

84. N. Saitou, M. Nei, The neighbor-joining method: a new method for reconstructing phylogenetic trees. Molecular biology and evolution 4, 406–425 (1987).

85. K. Tamura, G. Stecher, D. Peterson, A. Filipski, S. Kumar, MEGA6: Molecular Evolutionary Genetics Analysis version 6.0. Molecular biology and evolution 30, 2725–2729 (2013).

86. K. Tunyasuvunakool, J. Adler, Highly accurate protein structure prediction for the human proteome. 596, 590–596 (2021).

87. J. Mistry et al., Pfam: The protein families database in 2021. Nucleic Acids Research 49, D412–d419 (2021).

88. J. Jumper, R. Evans, A. Pritzel, T. Green, Highly accurate protein structure prediction with AlphaFold. 596, 583–589 (2021).

89. E. F. Pettersen et al., UCSF Chimera--a visualization system for exploratory research and analysis. Journal of Computational Chemistry 25, 1605–1612 (2004).

90. S. Jo, T. Kim, W. Im, Automated builder and database of protein/membrane complexes for molecular dynamics simulations. PloS one 2, e880 (2007).

91. J. Huang, S. Rauscher, G. Nawrocki, T. Ran, M. Feig, CHARMM36m: an improved force field for folded and intrinsically disordered proteins. 14, 71–73 (2017).

92. S. Jo, T. Kim, V. G. Iyer, W. Im, CHARMM-GUI: a web-based graphical user interface for CHARMM. Journal of Computational Chemistry 29, 1859–1865 (2008).

93. W. Humphrey, A. Dalke, K. Schulten, VMD: visual molecular dynamics. Journal of molecular graphics 14, 33–38, 27-38 (1996).

94. J. C. Phillips, D. J. Hardy, Scalable molecular dynamics on CPU and GPU architectures with NAMD. 153, 044130 (2020).

95. A. Waterhouse et al., SWISS-MODEL: homology modelling of protein structures and complexes. Nucleic Acids Research 46, W296–w303 (2018).

96. I. N. Shindyalov, P. E. Bourne, Protein structure alignment by incremental combinatorial extension (CE) of the optimal path. Protein Engineering 11, 739–747 (1998).

97. Schrödinger, The PyMOL Molecular Graphics System, Version 2.5. Schrödinger, LLC, New York, USA. (2020).

98. M. Mirdita, K. Schütze, ColabFold: making protein folding accessible to all. 19, 679–682 (2022).

99. J. Yang, Y. Zhang, I-TASSER server: new development for protein structure and function predictions. Nucleic Acids Research 43, W174–181 (2015).

100. E. Jurrus et al., Improvements to the APBS biomolecular solvation software suite. Protein science : a publication of the Protein Society 27, 112–128 (2018).

101. E. Smakowska-Luzan et al., An extracellular network of Arabidopsis leucine-rich repeat receptor kinases. Nature 553, 342–346 (2018).

102. Soejima Y. et al., Comparison of signal peptides for efficient protein secretion in the baculovirus-silkworm system. Open Life Sciences 8 (2013).

103. Y. Hashimoto, S. Zhang, S. Zhang, Y. R. Chen, G. W. Blissard, Correction: BTI-Tnao38, a new cell line derived from Trichoplusia ni, is permissive for AcMNPV infection and produces high levels of recombinant proteins. BMC Biotechnology 12, 12 (2012).

