## Supplementary material for "Arabidopsis Leucine Rich Repeat-Malectin Receptor Kinases in immunity triggered by cellulose and mixed-linked glucan oligosaccharides": Martin-Dacal_etal_SI_Figures_Datasets

#### *Appendix SI Information: Supplemental Figures*

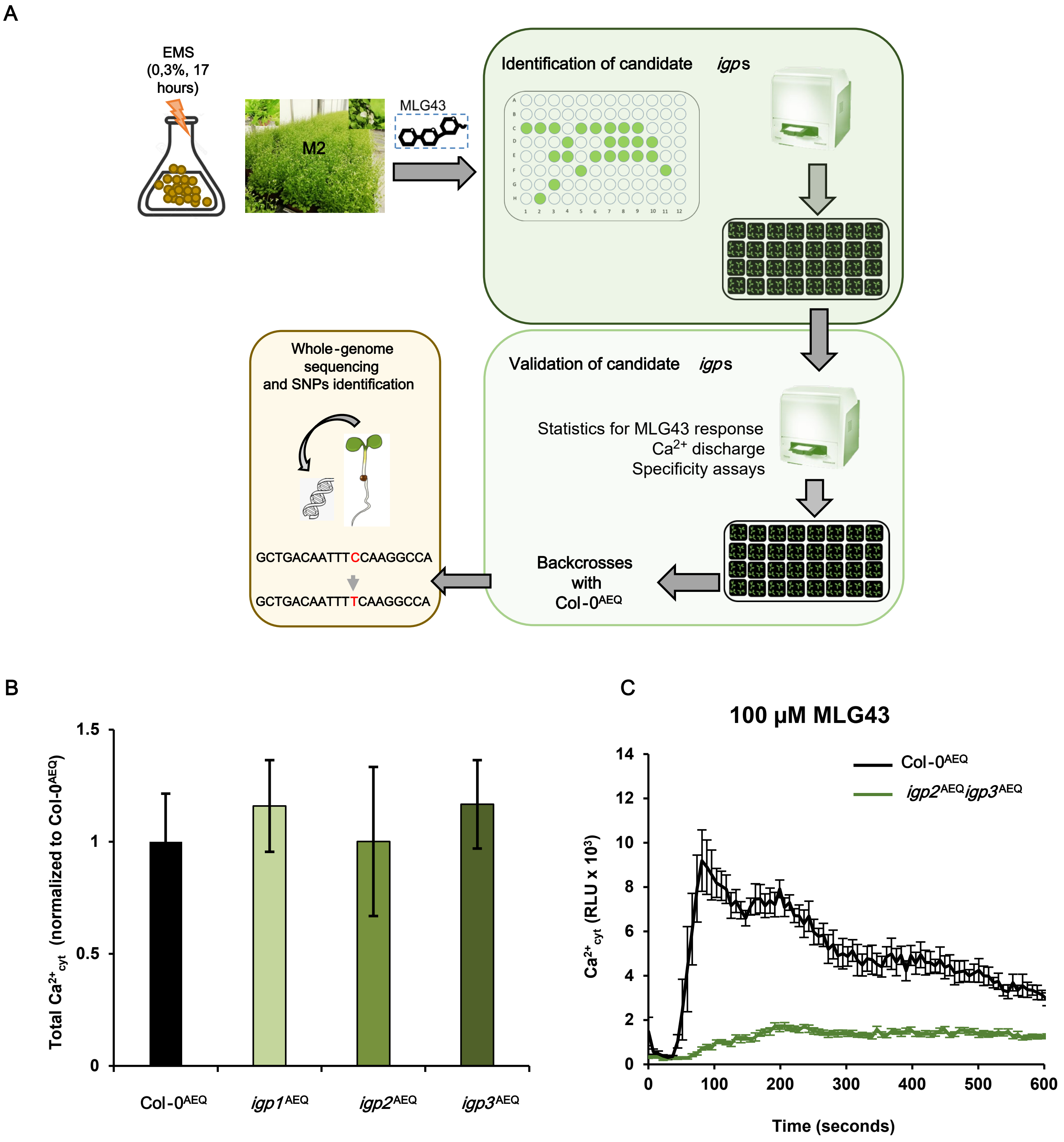


Supplemental figure 1

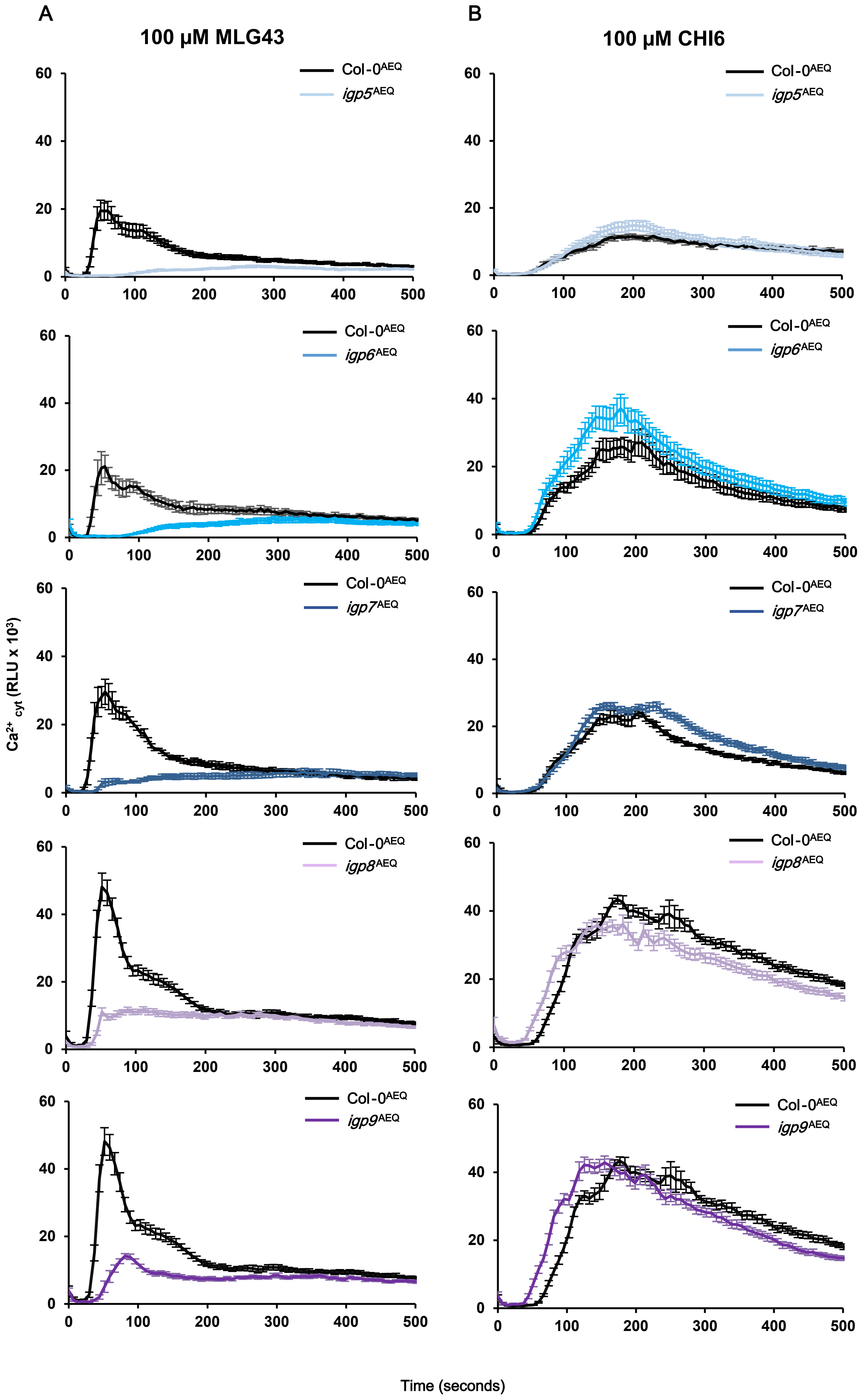


Supplemental figure 2

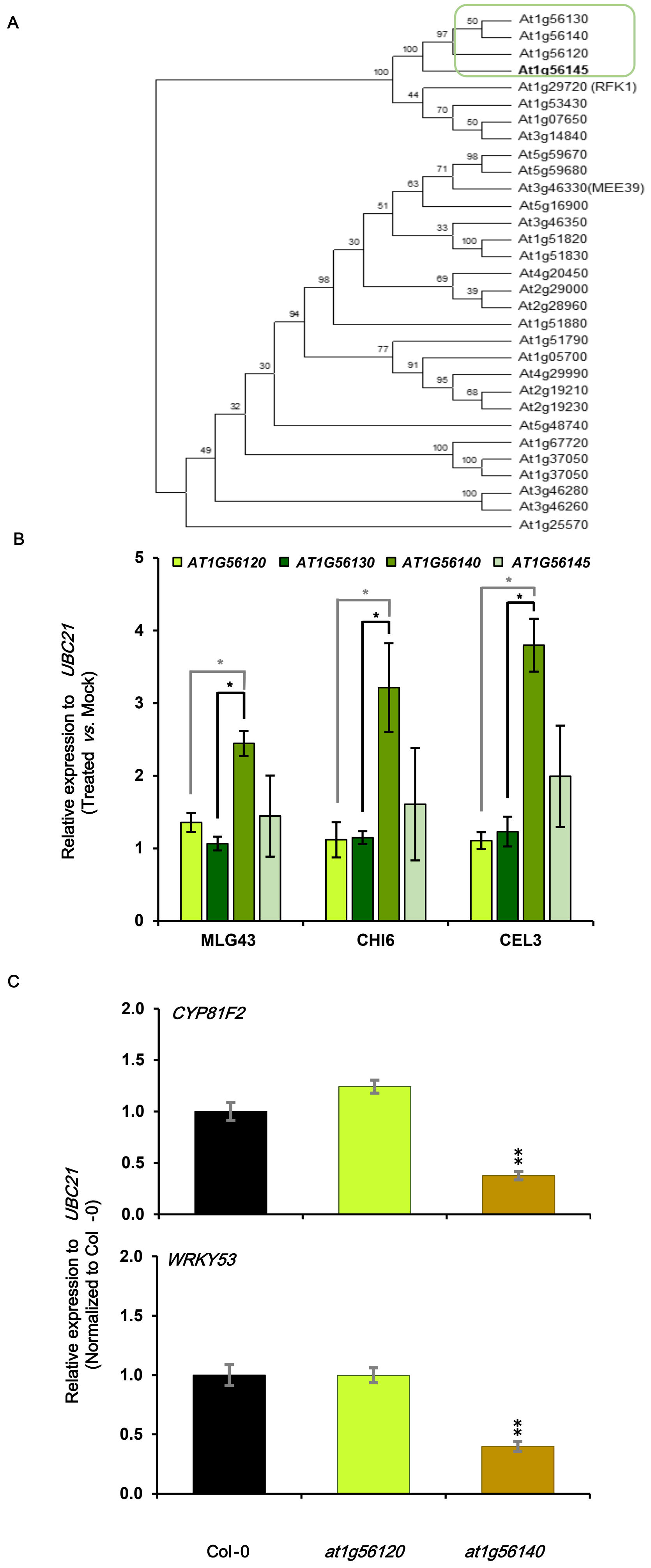


### Supplemental figure 3

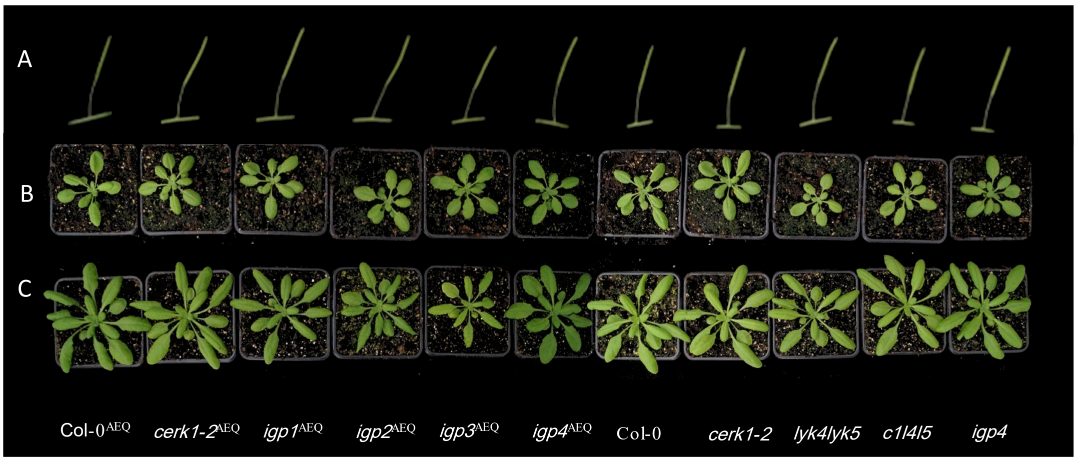


Supplemental figure 4

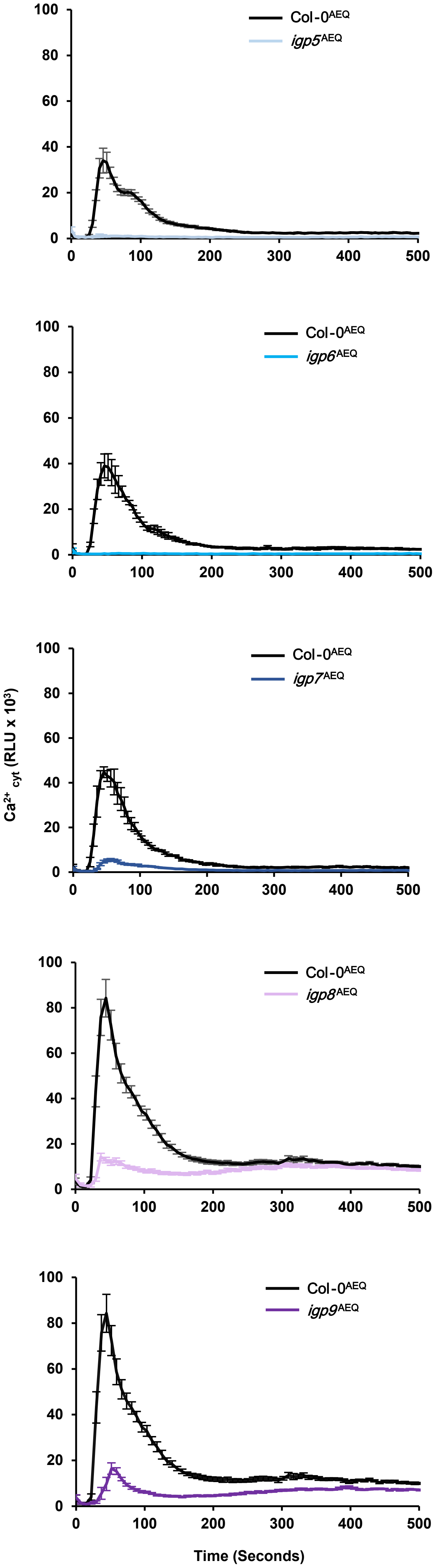


Supplemental figure 5

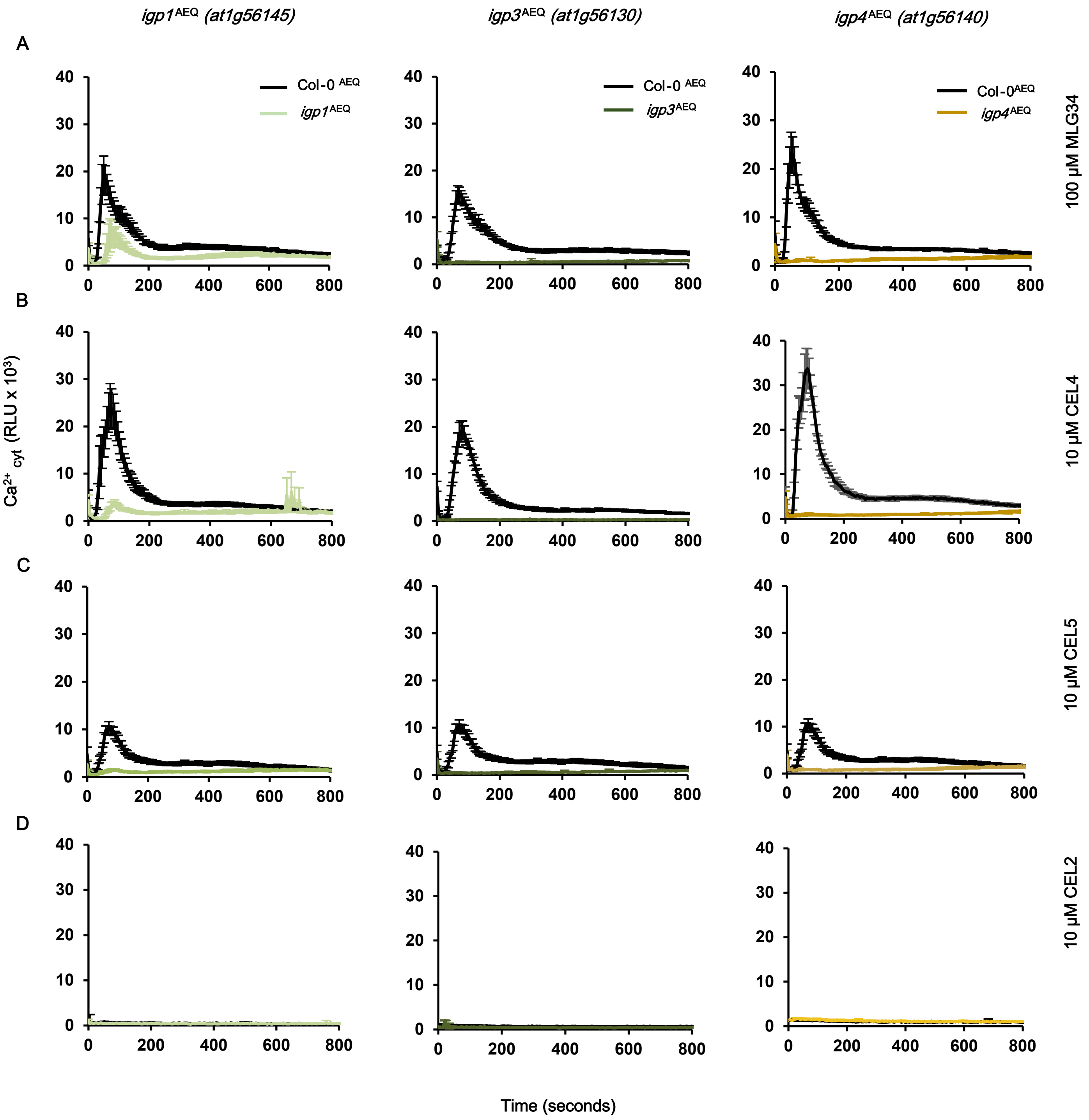


Supplemental figure 6

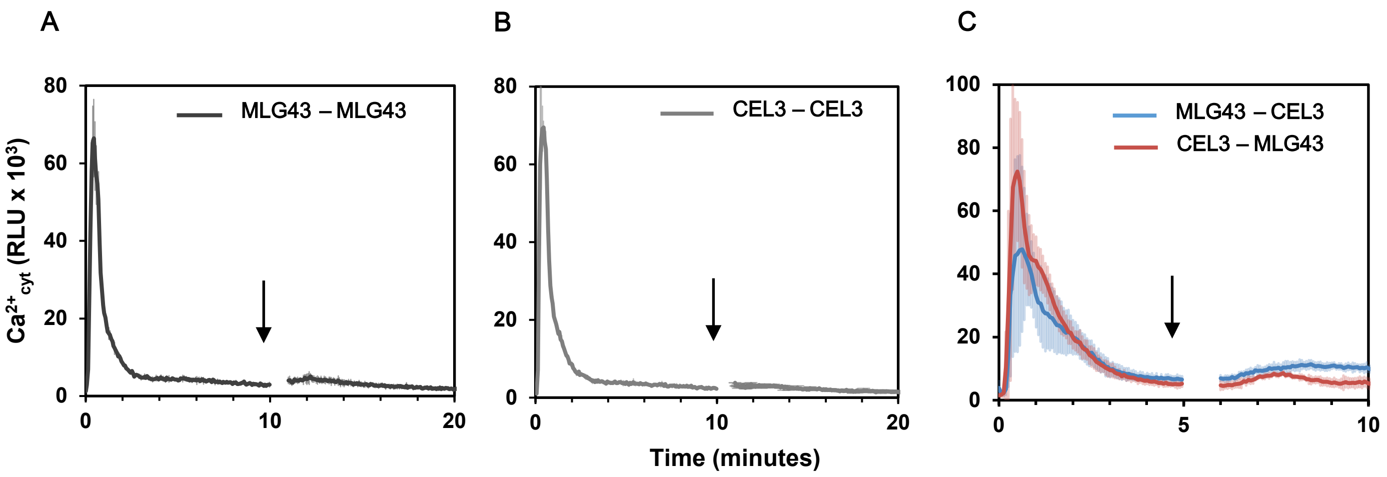


Supplemental figure 7

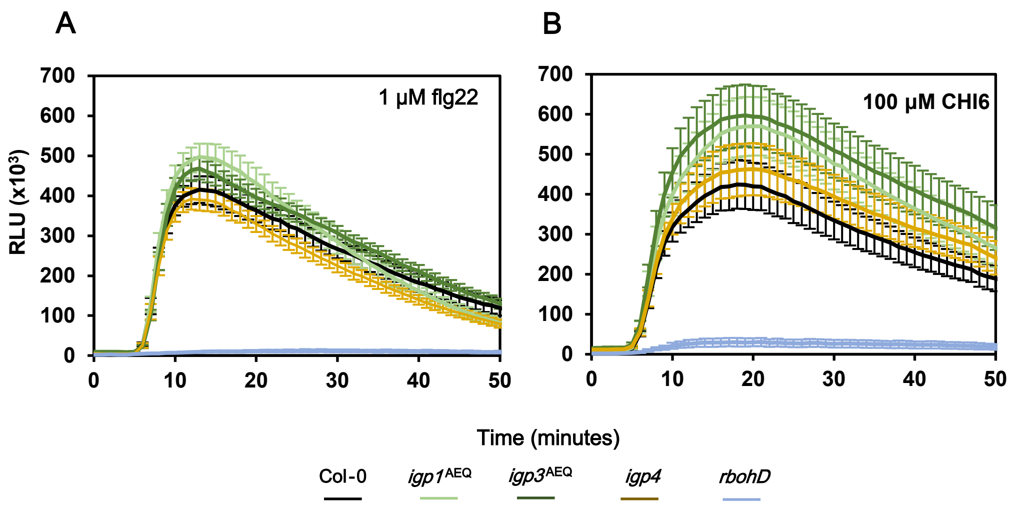


Supplemental figure 8

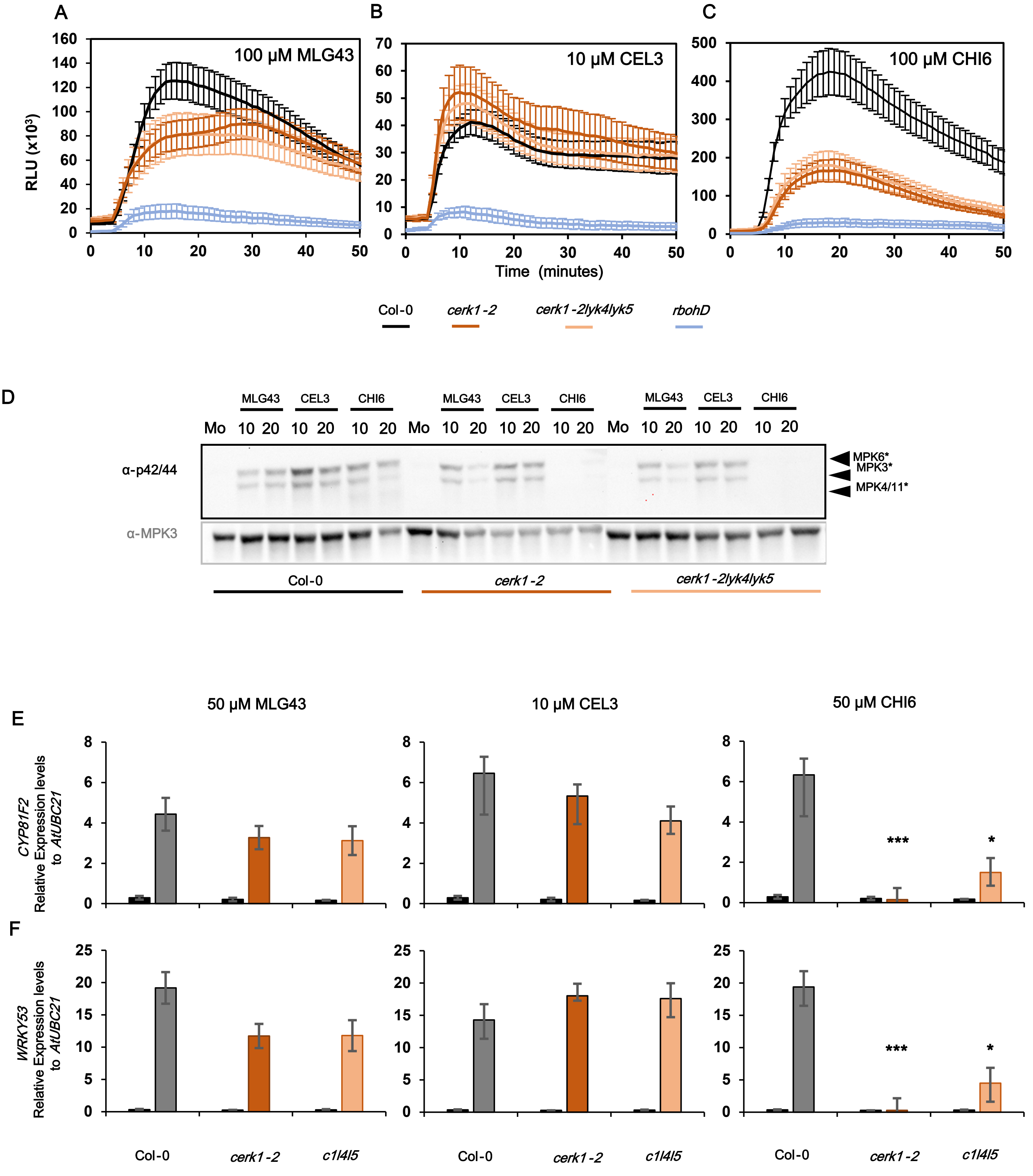


Supplemental figure 9

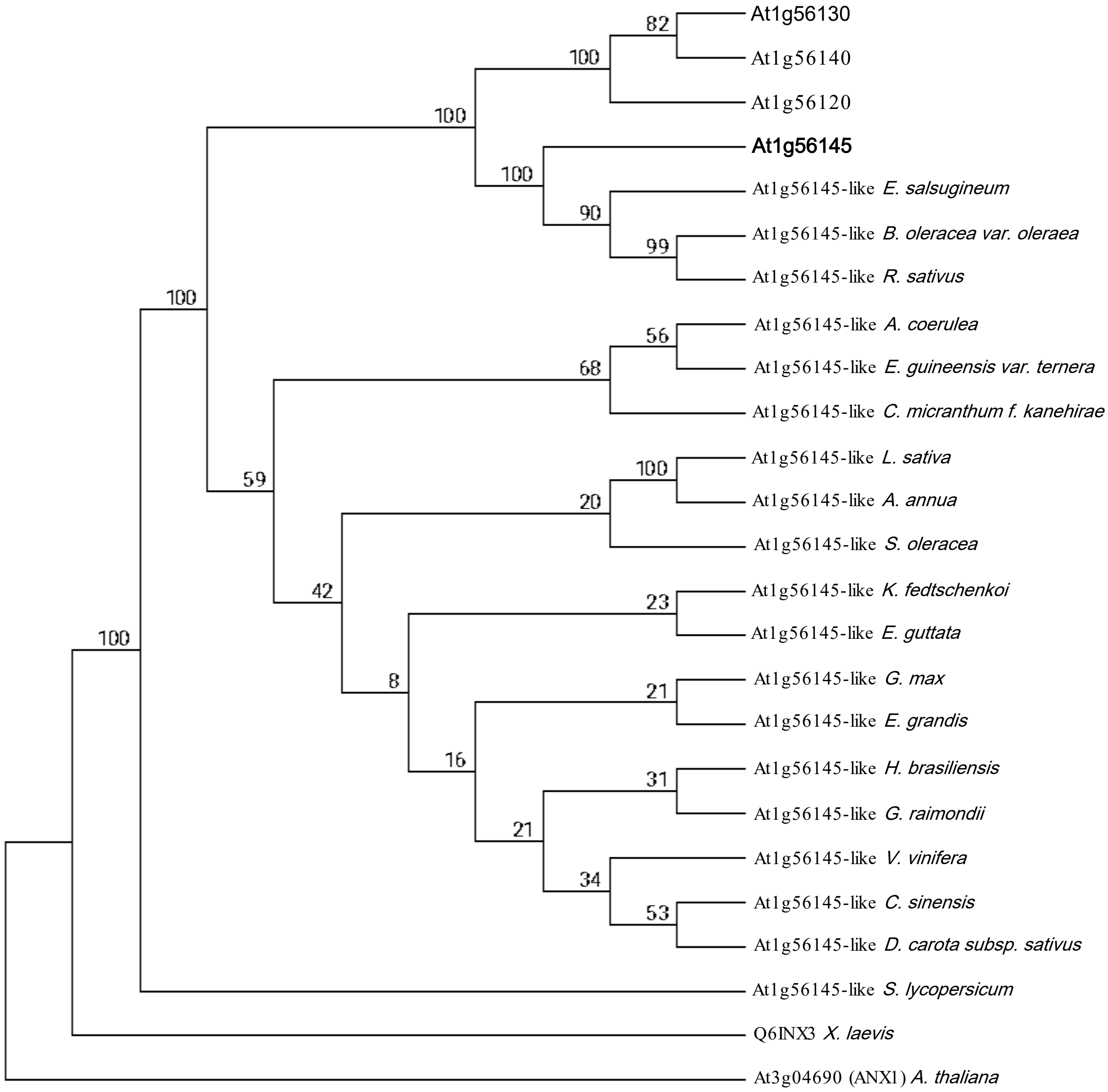


Supplemental figure 10

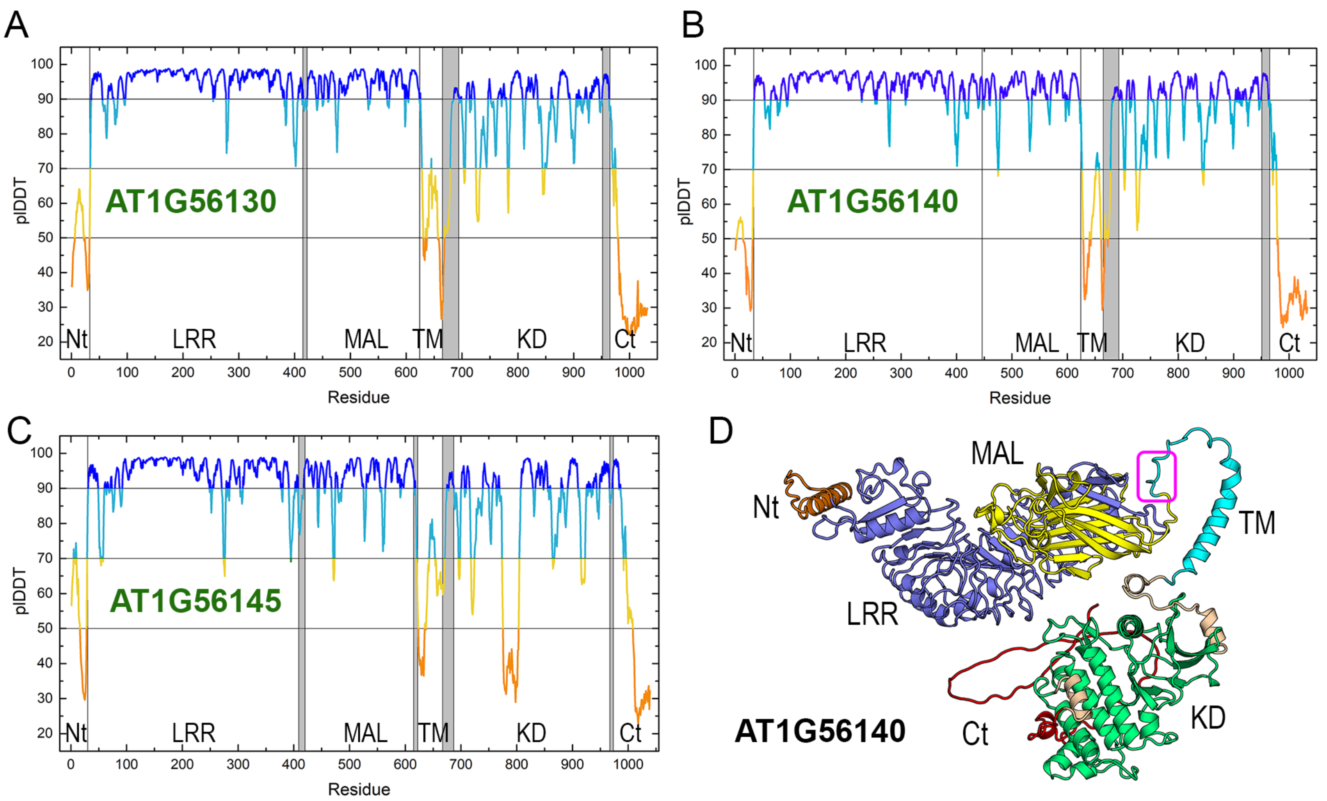


Supplemental figure 11

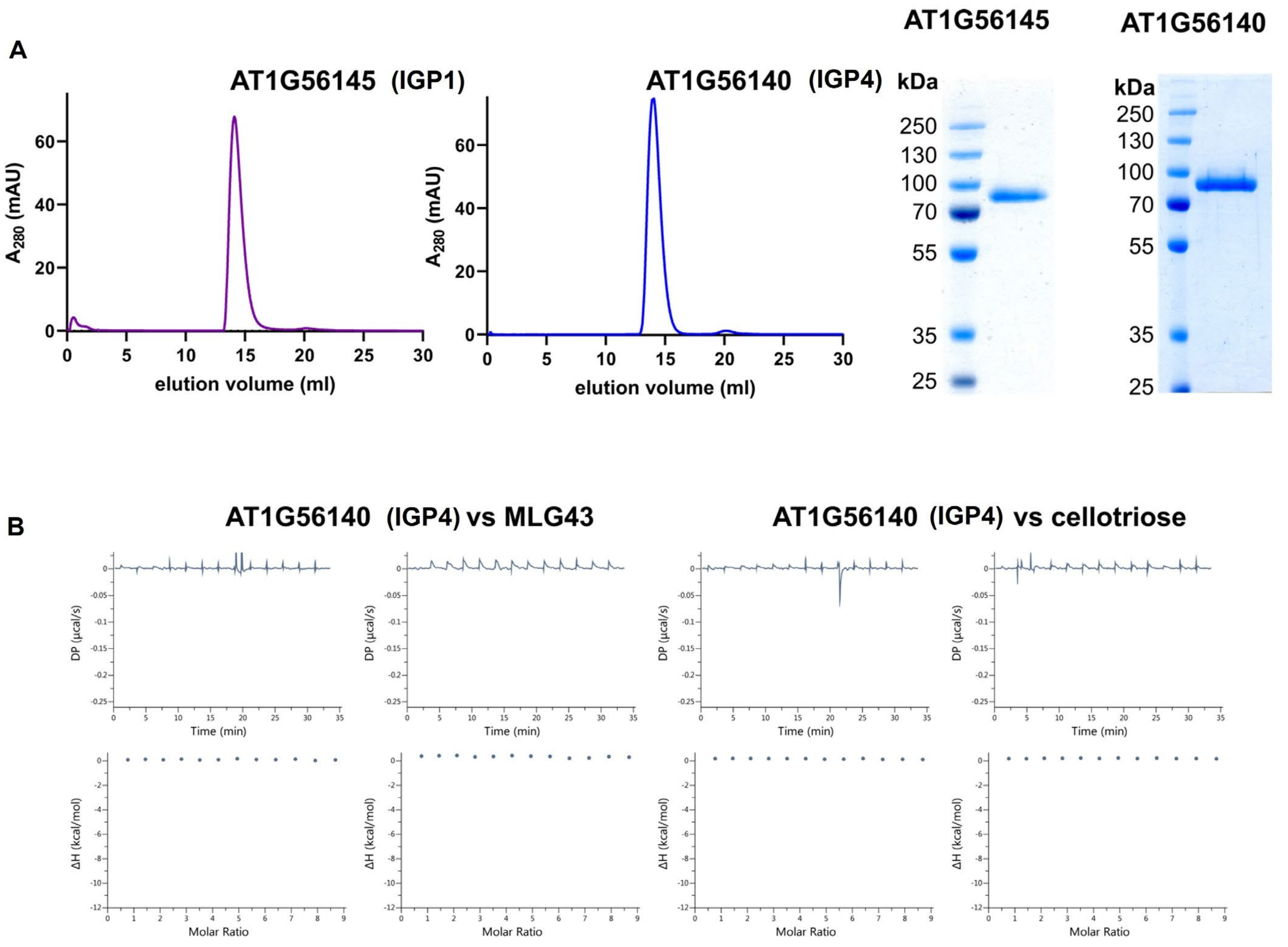


Supplemental figure 12

#### *Appendix SI Information: Datasets*

#### *SUPPORTING INFORMATION DATASETS*

**Dataset 1. NCBI whole genome sequence dataset:** The genome assembly data of *igp1*$^{AEQ}$, *igp2*$^{AEQ}$ *,* *igp3*$^{AEQ}$ , *igp4,* and Col-0$^{AEQ}$ can be retrieved from the NCBI Sequence Read Archive (SRA) under BioProject ID PRJNA864842 and Biosample accessions SAMN30087195, SAMN30087196, SAMN30087197, SAMN30087198, and SAMN30087199.

**Dataset 2. Chi-squared test statistics for *igp1*$^{AEQ}$-*igp3*$^{AEQ}$ mutant segregation.**

| | $H_0$ | *p*-value | Result |
|---|---|---|---|
| ***igp1***$^{AEQ}$ | recessive mutation | 0,7 > *P* > 0,5 | $H_0$ accepted |
| ***Igp2***$^{AEQ}$ | recessive mutation | 0,5 > *P* > 0,3 | $H_0$ accepted |
| ***Igp3***$^{AEQ}$ | recessive mutation | 0,8 > *P* > 0,7 | $H_0$ accepted |
| **Allelism *igp2*$^{AEQ}$ *x igp3*$^{AEQ}$** | allelic mutations | *P* > 0,95 | $H_0$ accepted |

#### Dataset 3. Chromosomal localization of SNPs mutations in *igp* mutants.

| Candidate | Locus | Position* | AF** | Protein | Change | Motif affected |
|---|---|---|---|---|---|---|
| *igp1* | ***AT1G56145*** | ***21008701*** | **0.99** | **LRR-MAL RLK** | **E to K (906)** | **Protein Kinase Domain** |
| *igp2* | ***AT1G56130*** | ***20995970*** | **1** | **LRR-MAL RLK** | **G to E (625)** | **Transferase(Phosphotransferase) domain1** |
| | *AT1G56140* | *21006273* | 1 | LRR-MAL RLK | 6th intron | |
| | *AT1G58050* | *21482658* | 1 | RNA helicase family protein | silent mutation | |
| | *AT1G57800* | *21412060* | 0.99 | ORTH3 | G to E (75) | Between zinc finger domains |
| *igp3* | ***AT1G56130*** | ***20995970*** | **1** | **LRR-MAL RLK** | **G to E (625)** | **Transferase(Phosphotransferase) domain1** |
| | *AT1G56140* | *21006273* | 1 | LRR-MAL RLK | 6th intron | |
| | *AT1G58050* | *21482658* | 0.99 | RNA helicase family protein | silent mutation | |
| | *AT1G58037* | *21474508* | 0.99 | Cysteine/Histidine-rich C1 domain family protein | R to H (10) | Near N-terminal end |
| | *AT1G58100* | *21513013* | 0.99 | TCP DOMAIN PROTEIN 8 | silent mutation | |
| | *AT1G54610* | *20394863* | 0.99 | Protein kinase superfamily protein | 5th intron | |

*** Nucleotide changes from C to T (expected mutations in EMS mutagenesis). **AF ≥0.99**

#### Dataset 4. Oligonucleotides used in this work.

| Type | Gene | Locus | Forward oligonucleotide | Reverse oligonucleotide |
|---|---|---|---|---|
| QPCR | *UBQ21* | *AT5G25760* | GCTCTTATCAAAGGACCTTCGG | CGAACTTGAGGAGGTTGCAAAG |
| | *CYP81F2* | *AT5G57220* | TATTGTCCGCATGGTCACAGG | CCACTGTTGTCATTGATGTCCG |
| | *WRKY53* | *AT4G23810* | CACCAGAGTCAAACCAGCCATTA | CTTTACCATCATCAAGCCCATCGG |
| | *IGP1* | *AT1G56145* | TTGGTTTGTGTTCATGTCTGGT | TATCTTGTTCAACGCCCGAG |
| | *IGP3* | *AT1G56130* | TCTCTCAGCGGATTTCAAATCA | TTGGCTTAAAGTCGCTGTCTC |
| | *IGP4* | *AT1G56140* | TCAATCAGCGGCTTTCCATT | CTAGCGTCGTTGTTTCTCGG |
| | *At1g56120* | *AT1G56120* | TGTTGAATGCTATTGACTGGTGT | CCGCGTCCTCCTTTTCAAAT |
| Genotyping | GABI_096F09 (*cerk1-2*) | *AT3G14840* | AGAATATATCCACGAGCACACGGTTCCAG | GACGAAAAGAGAGTGGATAAAGCAACCAC |
| | WiscDsLox297300_01C (*lyk4*) | *AT2G23770* | CATTTTCATCCATCGATGGAC | TTCCCTTTCACAACAATCCTG |
| | SALK_131911C (*lyk5*) | *AT2G33580* | TTCTGGTCTCAACCACCGTAC | CAGAAACCTGAGAGACGGATG |
| | SALK_005808 (*igp4*) | *AT1G56140* | AATGATACAGGTAATCCCCCG | GTTTGAGTCGAGAACACTGGC |
| | GABI-KAT LB | | CCCATTTGGACGTGAATGTAGACAC | |
| | SALK LBa1 | | TGGTTCACGTAGTGGGCCATCG | |
| | WiscDsLox LP | | AACGTCCGCAATGTGTTATTAAGTTGTC | |
